# Individuation of parts of a single object and multiple distinct objects relies on a common neural mechanism in inferior intraparietal sulcus

**DOI:** 10.1101/722256

**Authors:** Moritz F. Wurm, Katharine B. Porter, Alfonso Caramazza

**Affiliations:** Center for Mind/Brain Sciences, University of Trento, 38068 Rovereto (TN), Italy; Cognitive Neuropsychology Laboratory, Harvard University, Cambridge, MA 02138, USA

**Author notes:** Corresponding Author: Moritz F. Wurm, Center for Mind/Brain Sciences (CIMeC), University of Trento, Corso Bettini 31, 38068 Rovereto (TN), Italy,. MFW and KBP contributed equally to this work.

## Abstract

Object identification and enumeration rely on the ability to distinguish, or individuate, objects from the background. Does multiple object individuation operate only over bounded, separable objects or does it operate equally over connected features within a single object? While previous fMRI experiments suggest that connectedness affects the processing and enumeration of objects, recent behavioral and EEG studies demonstrated that parallel individuation occurs over both object parts and distinct objects. However, it is unclear whether individuation of object parts and distinct objects relies on a common or independent neural mechanisms. Using fMRI-based multivariate pattern analyses, we here demonstrate that activity patterns in inferior and superior intraparietal sulci (IPS) encode numerosity independently of whether the individuated items are connected parts of a single object or distinct objects. Lateral occipital cortex is more sensitive to perceptual aspects of the two stimulus types and the targets of the stimuli, suggesting a dissociation between ventral and dorsal areas in representing perceptual object properties and more general information about numerosity, respectively. Our results suggest that objecthood is not a necessary prerequisite for parallel individuation in IPS. Rather, our results point toward a common individuation mechanism that selects targets over a flexible object hierarchy, independently of whether the targets are distinct separable objects or parts of a single object.

## 1. Introduction

A fundamental capacity of human cognition is the ability to distinguish objects from the background, and to select a subset of objects for further processing, such as enumeration (e.g., counting apples in a bowl). Many theories support the differentiation of two stages in object processing: The first stage is object individuation, consisting of the selection of items at a particular location without specific knowledge of their features, which is followed by object identification, when a feature-rich representation becomes accessible (Kahneman, Treisman, & Gibbs, 1992; Leslie, Xu, Tremoulet, & Scholl, 1998; Pylyshyn, 1989; Xu, 2009; Xu & Chun, 2009). The individuation stage has the unique characteristic of being able to select multiple items in parallel without a behavioral cost. This has been demonstrated in the ability to track multiple moving targets at once (Howe, Cohen, Pinto, & Horowitz, 2010; Pylyshyn & Storm, 1988), as well as in the rapid enumeration of targets (Trick & Pylyshyn, 1994). Lesion and fMRI data also suggest that individuation and identification depend on and are processed in different cortical space, supporting their distinct roles in visual processing (Mishkin, Ungerleider, & Macko, 1983; Xu, 2009; Xu & Chun, 2006).

Parallel individuation has been studied extensively using behavioral methods. In the context of an enumeration task, parallel individuation is characterized by near zero errors, and short reaction times (RTs). Performance on these two measures changes dramatically based on the target set size. Within the ‘subitizing range’ (∼1-4), errors and RTs show little increase for each additional target, whereas there is a much larger cost for each additional target in the ‘counting range’ (∼5+) (Kaufman, Lord, Reese, & Volkmann, 1949). The different response profiles across these two ranges creates an ‘elbow’ as the slopes in error rates or RT change at ∼4 items, depending on the individual’s subitizing range (Akin & Chase, 1978). Thus, there is a limit to how many items can be individuated in parallel, which can be defined behaviorally for each individual.

Neural measures have also supported the existence of a limited capacity individuation mechanism. Recent work found that the N2pc, a parieto-occipital component assumed to reflect the operation of a capacity-limited attention-based individuation mechanism (Mazza & Caramazza, 2015), was modulated by target numerosity regardless of distracter presence, plateauing at 3 items (Ester, Drew, Klee, Vogel, & Awh, 2012; Mazza & Caramazza, 2011; Mazza, Pagano, & Caramazza, 2013). The location of the plateau was found to correlate with individual behavioral limits (Pagano, Lombardi, & Mazza, 2014; Pagano & Mazza, 2012) and is believed to relate to the precise individuation of items for further processing. Similar results have been observed in multiple-object tracking, again with modulation of the N2pc by target numerosity (Drew & Vogel, 2008).

In cortical space, fMRI studies have targeted the parietal lobe as the probable locus of this limited capacity individuation mechanism (Mitchell & Cusack, 2008; Todd & Marois, 2004). Activity in the inferior intraparietal sulcus (inferior IPS) is modulated by the number of objects existing in unique locations regardless of their features (Xu, 2009), and has a limit of about four objects (Todd & Marois, 2004; Xu & Chun, 2006). This is in contrast to the superior intraparietal cortex (superior IPS), where activity is modulated only by the number of objects with different features suggesting it is involved in object identification (Xu, 2009). Studies on number also converge on the parietal lobe, demonstrating activity modulated by target set size in neural regions also involved in spatial relations (Hubbard, Piazza, Pinel, & Dehaene, 2005). This overlap is consistent with an individuation mechanism tracking the number of items and their locations.

Previously, the large majority of functional neuroimaging experiments using an enumeration paradigm (Ansari, Lyons, van Eimeren, & Xu, 2007; Cutini, Scatturin, Basso Moro, & Zorzi, 2014; Damarla, Cherkassky, & Just, 2016; Knops, Piazza, Sengupta, Eger, & Melcher, 2014; Sathian et al., 1999) and studying parallel individuation (Xu, 2009; Xu & Chun, 2006) have asked subjects to individuate separate, unconnected target items. However, connectedness^1^ plays an important role when segmenting the visual field into separate units, and has been proposed to be the cue resulting in the initial organization of the visual field (Palmer & Rock, 1994). Connectedness also plays an important role in the allocation of attention (Driver & Vogel, 1998), and has been demonstrated to aid or interfere with different tasks. Balint’s syndrome is an example where connectedness aids performance. Whereas patients suffering from Balint’s syndrome are normally only able to visually apprehend a single object, adding a connecting line between two objects allows their attention to spread to include both objects and successfully compare the features of the two items (Humphreys & Riddoch, 1993). Connectedness has been shown to affect parallel individuation negatively in a behavioral multiple object tracking task with typical participants. When target stimuli were manipulated so that they were perceived as either connected to a distracting element or independently moving and disconnected from any distractors, performance suffered in the connected conditions (Scholl, Pylyshyn, & Feldman, 2001).

Whether connectedness has a negative effect on parallel individuation is still under debate. Recent behavioral work demonstrated the presence of subitizing for the enumeration of both multiple separate objects and connected parts of a single object (Porter, Mazza, Garofalo, & Caramazza, 2016). Likewise, a recent EEG study found that the N2pc was modulated as a function of number of connected object parts within the subitizing range in a similar way as it would be expected for unconnected, bounded targets (Poncet, Caramazza, & Mazza, 2016). These findings invite the inference that a common neural mechanism may underlie the individuation of object parts and multiple objects. However, this interpretation seems at variance with prior studies indicating an effect of connectedness on individuation performance (Fornaciai, Cicchini, & Burr, 2016; Franconeri, Bemis, & Alvarez, 2009; He, Zhang, Zhou, & Chen, 2009; He, Zhou, Zhou, He, & Chen, 2015; Scholl et al., 2001). Neuroimaging studies have also suggested that connectedness could affect the individuation of targets. An fMRI adaptation study found that connecting target dots with lines in an enumeration task caused a shift in the adaptation curves in IPS, suggesting a decrease in the encoded numerosity for those displays (He et al., 2015). Another study investigated the effects of grouping on neural activity in the inferior IPS, and reported that grouped displays resulted in lower levels of activity than ungrouped displays (Xu & Chun, 2007). These studies suggest that individuation of parts of a single object and multiple objects relies on distinct neural mechanisms. However, while He at al. (2015) found effects of connectedness within the subitizing range behaviorally, they only investigated neural effects of connectedness for 5+ targets. Xu & Chun (2007) also found neural effects of connectedness but did not manipulate numerosity. Taken together, the current neuroimaging evidence cannot disambiguate whether individuation of parts of a single object and multiple objects underlies a common or two independent neural mechanisms.

In the current fMRI experiment, we investigate the neural effects of connectedness on individuation in the parietal cortex, including set sizes within the range of parallel individuation. Specifically, the study is designed to test whether neural response patterns associated with numerosities in the subitizing range are similar for, and even generalize across, multi-object and single-object stimuli with multiple objects and connected parts of a single object (Figure 1). If the IPS is sensitive to connectedness, as is hinted by He et al. (2015) and Xu & Chun (2007), then we expect differential response patterns to connected and unconnected sets of targets. The most extreme outcome would be that there would be no modulation by target number for the connected items, as all set sizes are viewed as one object. In contrast, if the role of the inferior IPS were to individuate target items across different frames of reference and definitions of figure and ground, then we would predict similar modulation by set size for both independent objects and connected targets of a single object. A previous study has shown some evidence of cross-classification of number across tasks in the posterior parietal cortex, indicating there is some ability to generalize number across stimulus types (Knops et al., 2014). Consequently, we also expect to see above-chance classification of numerosity in the IPS across stimulus type if individuation of parts of a single object and multiple objects relies on a common neural mechanism. An additional aspect concerns the role of visual areas in processing numerosity at perceptual vs. more abstract levels of representation. Recent studies suggest that ventral visual areas are involved in representing perceptual aspects of numerosity independently of task (Cavdaroglu, Katz, & Knops, 2015; DeWind, Park, Woldorff, & Brannon, 2019; Fornaciai & Park, 2018). While parietal cortex might represent numerosity independently of connectedness due to explicit task demands, it is unclear whether, and if so, how visual areas represent numerosity, in particular with regard to the single object displays which are ambiguous with regard to numerosity: does visual cortex represent them as single objects (a circle) independently of the number of arcs or is visual cortex also sensitive to the number of target numerosity across the different stimulus displays? Therefore, in addition to IPS, we also investigate neural responses to target individuation in the lateral occipital complex (LOC), a region involved in object shape processing.

**Figure 1.**
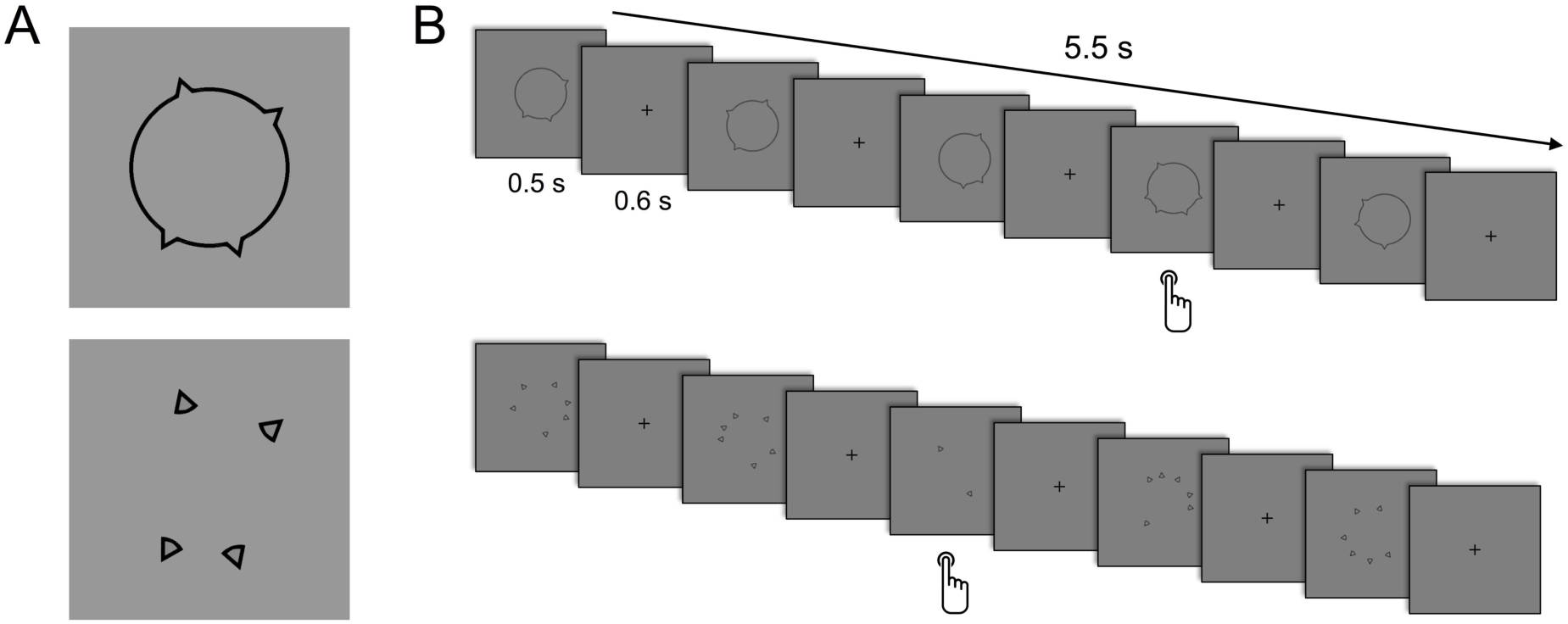
Stimuli and experimental design. (A) An example of the matched stimuli for the two stimulus types. Note that the unconnected arcs in the multi-object display (bottom) were jittered slightly in their rotation to avoid the perception of an illusory circle. (B) Time sequence of a block for the main experiment. Examples depict oddball blocks, with the expected response point, for each stimulus type (top: single-object displays, bottom: multi-object displays).

## 2. Material and Methods

We report how we determined our sample size, all data exclusions, all inclusion/exclusion criteria, whether inclusion/exclusion criteria were established prior to data analysis, all manipulations, and all measures in the study

### 2.1 Participants

14 participants between the ages of 20 and 26 (10 female) were scanned at the University of Regensburg and compensated for their participation. The sample size was determined based on previous studies using comparable parallel individuation paradigms (Xu & Chun, 2006, 2009). One participant was excluded from analyses due to incomplete data collection as a result of computer failure. All participants were right-handed and had normal or corrected-to-normal vision. The study was approved by the Committee on the Use of Human Subjects in Research at Harvard University, the Ethics Committee for Experimentation with Human Beings at the University of Trento, and the Faculty of Medicine Ethics Committee at the University of Regensburg.

### 2.2 Stimuli and Design

There were 12 stimulus conditions (6 numerosities x 2 stimulus types). Numerosities were from 1 to 6; the two stimulus types were single-object displays with connected targets and multi-object with unconnected targets. The targets of single-object displays consisted of a black outline of a circle with protruding arcs. multi-object displays consisted of outline-closed arcs (Figure 1A). The locations of the single-object display target features were generated randomly for each stimulus and each participant, constrained such that no two features could overlap. The orientation of each arc in the multi-object displays was randomly selected from a set of angles (−20°, -10°, 0°, +10°, +20°) to avoid the illusory percept of a circle. The goal of the use of outline stimuli instead of filled shapes was twofold: to eliminate any possible percept of the protruding arcs as separate shapes occluding the central circle in the single-object condition, and to better match the low-level visual properties of the two stimulus types. A unique stimulus set was created in advance and presented for each participant using Matlab with the Psychophysics Toolbox extensions (Brainard, 1997; Kleiner et al., 2007; Pelli, 1997); no stimulus was presented twice. However, the locations of the targets were controlled across stimulus types, such that for each unique grouped stimulus, a matched ungrouped stimulus also existed (Figure 1A). Data from a previous study (Porter et al., 2016) demonstrated that the grouped and ungrouped outline stimuli showed behavioral evidence of parallel individuation.

Each run was presented as a mini-block design, with 5 stimulus presentations per block. Subjects performed an oddball detection task: They were instructed to observe the sequentially presented stimuli and to press a button when the numerosity of a stimulus was different from that established by the first stimulus in the block. Each block was 5.5 s long, with 500 ms presentation, 600 ms inter-stimulus-interval, followed by a 2.5 second inter-block-interval, resulting in 8 seconds between the onset of each block (Figure 1B).

Forty-seven blocks were presented per run: 36 ‘pure’ blocks without oddball stimuli (3 repetitions x 6 numerosities x 2 stimulus types), 7 blocks containing an oddball trial, and 4 fixation blocks (8 s long). An additional 8 seconds of fixation was added at the beginning and end of each run. This organization resulted in fixation for 12.2% of the run, and 14.9% of the blocks including an oddball trial. Six runs were collected for each participant, resulting in 18 blocks per stimulus condition. Since the number of oddball blocks did not divide evenly into the number of stimulus types, we pseudorandomly determined the stimulus type in the following manner: over the 42 possible oddball blocks in all 6 of the runs, 3 repetitions occurred for each stimulus condition; the remaining 6 blocks were split into 1 block per numerosity, with the stimulus type (grouped, ungrouped) determined randomly for each block. These oddball blocks were excluded from all analyses.

The order of the blocks in each run was pseudorandomly determined in the following manner for each subject. First, the position of the 4 fixation blocks and 7 oddball blocks were determined randomly, with the constraint that fixation could not occur in the first or last block of the run. The pure stimulus blocks were then inserted into the remaining available positions: the twelve stimulus conditions were randomly shuffled, and then inserted into the first 12 available slots. This was repeated two more times to fill the remaining block positions.

### 2.3 Scanning Parameters

Functional and anatomical data were collected using a 3-Tesla Allegra scanner (Siemens, Erlangen, Germany) at the University of Regensburg with a single channel head coil. Functional data were collected using ascending interleaved slice acquisition with a T2*-weighted gradient echo planar imagine (EPI) sequence (image matrix: 64 x 64 with 34 axial slices, repetition time = 2000 ms, echo time (TE) = 30 ms; flip angle (FA) = 90°, field of view (FOV) = 192 x 192 mm^2^, slice thickness = 3 mm, gap = 0.30 mm, with 3×3 mm in-plane resolution). Structural data were acquired using a high-resolution scan (160 sagittal slices, 1×1×1 mm^3^) with a T1-weighted MP-RAGE sequence (TR = 2.25 s, TE 2.6 ms, FA = 9°, FOV = 240 x 256 mm). The sequence was optimized for the differentiation of gray and white matter by using parameters from the Alzheimer’s Disease Neuroimaging Initiative project (http://adni.loni.ucla.edu/). Stimuli were generated using Psychophysics toolbox for MATLAB and projected onto a screen inside the bore with a LCD projector (JVC, DLA-G20, Yokohama, Japan), which was viewed by a mirror attached to the head coil.

### 2.4 Preprocessing

The fMRI data were analyzed using SPM8, CoSMoMVPA (Oosterhof, Connolly, & Haxby, 2016), as well as custom Matlab software. The first 4 volumes were removed to avoid T1 saturation. The first volume of the first run was aligned to the high-resolution anatomy (6 parameters). Data were 3D motion corrected (trilinear interpolation, with the first volume of the first run of each participant as reference), followed by slice time correction, and high-pass filtering (cutoff frequency of 3 cycles per run). Spatial smoothing was applied with a Gaussian kernel of 6 mm FWHM. Anatomical and functional data were transformed into MNI space using trilinear interpolation.

### 2.5 Regions of Interest

Regions of interests (ROIs) for LOC, inferior IPS, and superior IPS were defined using functional localizer coordinates reported in Xu (2009; MNI coordinates, converted from Talairach space, x/y/z, left LOC: -36/-81/2, right LOTC: 47/-58/14, left inferior IPS: -30/-86/27, right inferior IPS: 28/-82/28, left superior IPS: -24/-64/40, right superior IPS: 25/-64/47). For univariate analyses, a 5.5 mm radius sphere was built around the coordinate voxel, resulting in ∼27 voxels selected, to approximate the same size as previously used in the literature (Xu, 2009). For multivariate analyses, a larger ROI size of 9 mm radius sphere was used to increase the number of features for classification, resulting in ∼122 voxels selected. ROIs were averaged across hemispheres.

We also performed ROI analyses using individually defined ROIs based on our own functional localizer experiments (visual short-term memory task for superior IPS, object localizer for LOC and inferior IPS) similar to those reported by Xu (2009). However, a comparison of these ROIs (MNI coordinates, x/y/z; left LOC: -30/-83/20, right LOC: 37/-77/18, left inferior IPS: -26/-75/36, right inferior IPS: 27/-73/38, left superior IPS: -26/-62/47, right superior IPS: 31/-59/51) with those identified by Xu revealed satisfactory overlap only in superior IPS (distance between peaks, left/right: 7.5/7.3 mm), whereas inferior IPS and LOC deviated considerably from Xu’s ROIs (inferior IPS: 14.8/13.5 mm distance; LOC: 19.1/21.8 mm distance), which might indicate a mislocalization of the latter areas (Supplementary Fig. 1A). We therefore decided to use coordinates by Xu (2009) for the ROI analyses reported here. However, results of the ROI analysis using individually defined ROIs can be found in Supplementary Figures 1-3. These analyses revealed no substantially different results.

### 2.6 Univariate analysis

For each participant and run, beta weights of the experimental conditions were estimated using design matrices containing predictors of the 12 conditions, oddball blocks, and of 6 parameters resulting from 3D motion correction (x, y, z translation and rotation). Each predictor was convolved with a dual-gamma hemodynamic impulse response function (Friston et al., 1998). Each block was modeled as an epoch lasting from block onset to offset (5.5 s). The resulting reference time courses were used to fit the signal time courses of each voxel. The resulting beta weights were averaged across the 6 runs to obtain one beta weight per condition and voxel.

We first investigated the effect of the different stimulus types on the level of activity in each ROI. For each participant, ROI, and condition, we extracted the betas from the ROI and averaged the beta values across voxels. We performed a 2 x 6 repeated-measures ANOVA to investigate potential main effects and interactions between stimulus type and numerosity. We corrected the statistical results for the number of ROIs (= 3) using the false discovery rate (FDR) at q = 0.05. To ensure that low-level factors were not driving our results, we replicated the GLM and analyses with number of black pixels in each image presented as a regressor of non-interest. To investigate the point at which activity plateaued in each region, we performed, for each ROI and stimulus type, paired two-tailed t-tests comparing beta coefficients between the largest three numerosities (4 vs. 5, 5 vs. 6), FDR-corrected for the number of ROIs, stimulus types, and tests per ROI (= 12). A plateau was assumed if there was no significant difference in activity between two numerosities. However, a lack of significance does not prove the null hypothesis that activity for two numerosities is at the same level. To further validate at which numerosity a plateau is reached, we additionally performed a Bayesian model comparison, which allowed comparing the likelihoods for the hypothesis that a plateau is reached against the alternative hypothesis that no plateau is reached. To this end, we computed Bayes factors for different linear regression models, using the lmBF function of the BayesFactor package (version 0.9.12) for R (version 3.4.2) (Morey, Rouder, Jamil, & Morey, 2015) and a default Cauchy prior width of r = 0.707 for effect size. Participants were included in the models as random factor. We tested three models: (1) a plateau is reached with 4 items (weights: 1,2,3,4,4,4), (2) a plateau is reached with 5 items (weights: 1,2,3,4,5,5), and (3) no plateau is reached within the tested range (i.e., a plateau is reached with 6 items or more; linearly increasing weights: 1,2,3,4,5,6). In a second step, we computed the relative Bayesian evidence as the ratio between the Bayes factors, which indicates how much more likely model X is relative to model Y. Likelihood ratios between Bayes factors > 3 indicate moderate (and noteworthy) evidence in favor of one model vs. the other (Jeffreys, 1961).

### 2.7 Multivoxel pattern analysis (MVPA)

We performed two kinds of multivoxel pattern analysis (MVPA): classification and representational similarity analysis. MVPA was performed in ROIs and in the whole brain using a searchlight with the same radius of 9 mm. Whole brain analyses were conducted to identify effects in regions other than the tested ROIs and to characterize anatomical location, extent, and peaks of information-coding clusters in more detail. No feature selection was used, i.e., all ∼122 voxels per ROI / searchlight sphere were used for MVPA.

For ROI analyses, to identify significant classification accuracies above chance, we corrected statistical results (one-sided one sample t tests) for the number of ROIs and tests/models (=12) using the FDR at q = 0.05. Statistical maps were corrected for multiple comparisons using a cluster-based Monte Carlo simulation algorithm as implemented in the CoSMoMVPA Toolbox (Oosterhof et al., 2016). We used a threshold of p = 0.05 at the cluster level, and initial voxelwise threshold of p = 0.001, and 10000 iterations of Monte Carlo simulations. For visualization, maps were projected on a cortex surface of a Colin27 MRI volume as provided by the Neuroelf toolbox using BrainVoyager QX 2.8 (BrainInnovation).

### 2.8 Classification analysis

For classification analysis, we used a linear discriminant analysis (LDA) classifier and beta maps for each run, condition, and participant as computed in the univariate analysis. Thus, each beta map was based on 3 blocks, and there were 6 beta maps per condition (18 beta maps per condition in total). To remove amplitude effects of the different conditions as much as possible, we demeaned the data for each condition and voxel pattern of a ROI / searchlight sphere by subtracting the mean beta of the voxel pattern from each beta of the individual voxels. First, we tested whether a given region encodes information about numerosity independent of stimulus type, i.e., whether representations of numerosity generalize across single-object and multi-object displays. To this end, we trained the classifier to discriminate the 6 numerosities of the single-object displays and tested the classifier on the multi-object displays, and vice versa. Numerosities were classified using multiclass decoding, i.e., each numerosity was discriminated from the remaining 5 numerosities, and resulting accuracies were averaged. Mean accuracies for each participant and searchlight/ROI sphere were entered into a one sample t test against chance. Chance for this classification was 16.7%, as there were 6 numerosities. Second, we investigated whether a given region encodes information about stimulus type independent of numerosity, i.e., whether representations of single-object and multi-object displays generalize across numerosity. To this end, we trained a classifier to discriminate the two stimulus types (single-object and multi-object displays) on 5 out of the 6 numerosities, and tested on the remaining numerosity. This was iterated 6 times, leaving out each numerosity once. Resulting accuracies were averaged across iterations and entered into a one sample t test against chance (50%).

### 2.9 Representational similarity analysis

To investigate the representational content of brain regions with regard to specific hypothesized models, we performed a multiple regression representational similarity analysis. To compute pairwise correlations between each of the 12 conditions, we used the beta maps of the univariate analysis that were averaged across runs. For each searchlight/ROI sphere, we extracted the mean beta values to obtain one multivoxel pattern per condition. Each voxel pattern was demeaned as in the classification analysis. Next, we correlated the patterns with each other resulting in a 12 x 12 correlation matrix per ROI/sphere and participant. Correlation matrices were inverted (1 – *r*) into representational dissimilarity matrices (RDMs).

The following model-based RDMs were used (Figure 4B): (1) The Object Individuation model assumes that representation of numerosity in IPS is sensitive to connectedness of targets. Thus, single-object displays, which have connected targets, are all represented as one object (i.e., numerosity = 1). For multi-object displays, which have unconnected targets, numerosities in the subitizing range (1-4) linearly increase while higher numerosities are not further differentiated (Figure 4A, left). The RDM was generated by computing the Euclidean distances between the numerosity representations associated with the 12 stimulus items. (2) The Feature Individuation model assumes that representation of numerosity in IPS is not altered by connectedness of items. Thus, for both single-object and multi-object displays, numerosities in the subitizing range (1-4) linearly increase while higher numerosities are not further differentiated (Figure 4A). (3) To ensure that putative effects in IPS are not merely artifacts of difficulty in looking for deviant numerosities in a block, we included Reaction Time and Accuracy models (Figure 4A, B). These models are expected to reflect how different the blocks are in terms of speed and accuracy to detect whether numerosities of trials are same or different (see *Behavioral Experiment* for details). (4) Single- and multi-object displays differ in terms of the spatial extent of the displays: Single-object displays with connected targets always occupy the same spatial area (a circle), whereas the spatial extent of the area in which the unconnected targets occur increases with increasing numerosity. To capture perceptual and attentional effects related to the processing of targets within areas of varying size, we computed Spatial Extent model of the area in which targets could occur (Figure 4A, B). For the multi-object displays we computed the spatial extent by adding up the side lengths of the convex hull formed by the targets. This was done for each possible combination of target locations, followed by averaging across combinations. For numerosity = 1, we defined a spatial extent of 0. For numerosity of 2, we computed the distance between target locations multiplied by 2. For the single-object displays, we computed the circumference of the circle (2pi * r), which was identical for all numerosities. (5) Finally, we included a Stimulus Type model that distinguishes single-object and multi-object displays in a categorical manner (Figure 4B).

**Figure 2.**
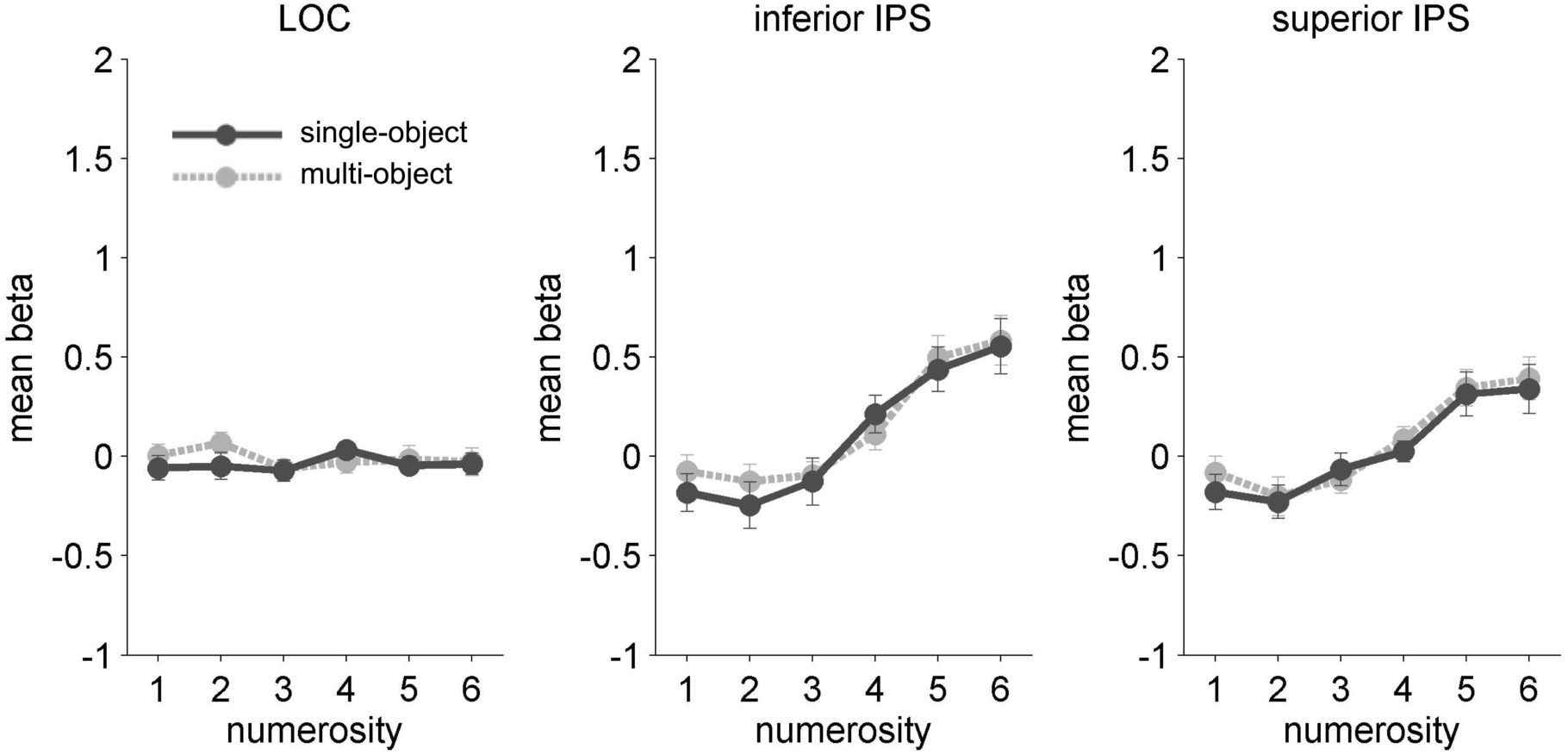
Univariate results. Mean beta coefficients in response to numerosity and stimulus type for each ROI. Error bars indicate SEM.

**Figure 3.**
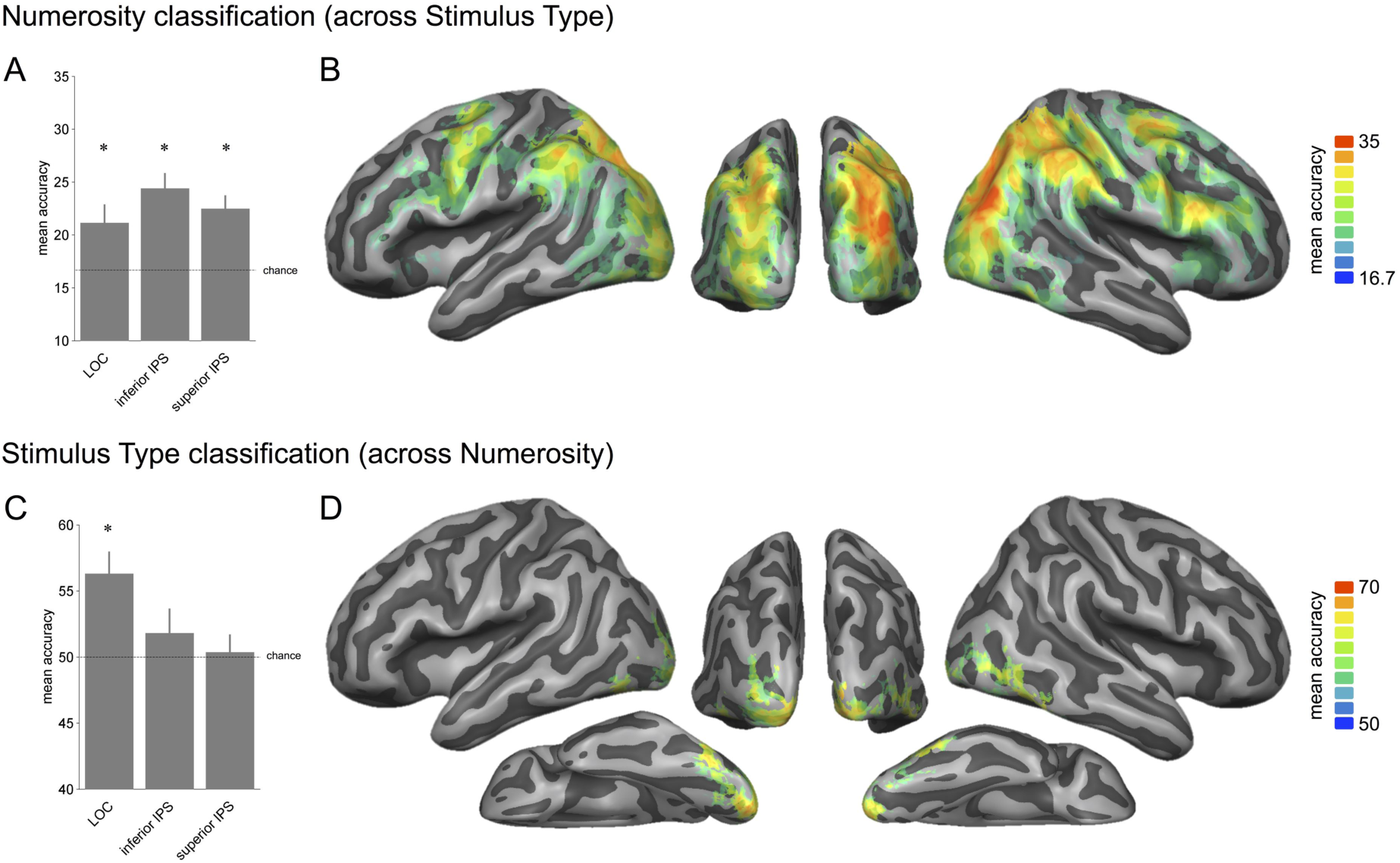
Classification of Numerosity and Stimulus Type. In Numerosity classification (A and B), the classifier was trained to discriminate numerosities in single-object displays and tested on its accuracy to discriminate numerosities in the multi-object displays (and vice versa). Accuracy for decoding at chance is 16.7%. In Stimulus Type classification (C and D), the classifier was trained to discriminate single-object vs. multi-object displays for four of five numerosities and tested on its accuracy to discriminate single-object vs. multi-object displays on the held out numerosity (iterated for all numerosities). Accuracy for decoding at chance is 50%. A and C show the results of the ROI analysis, with asterisks indicating FDR corrected significant above chance decoding. Error bars indicate SEM. B and D show mean accuracy maps of the searchlight analysis (cluster size corrected for multiple comparisons at p = 0.05, initial voxel threshold p = 0.001).

**Figure 4.**
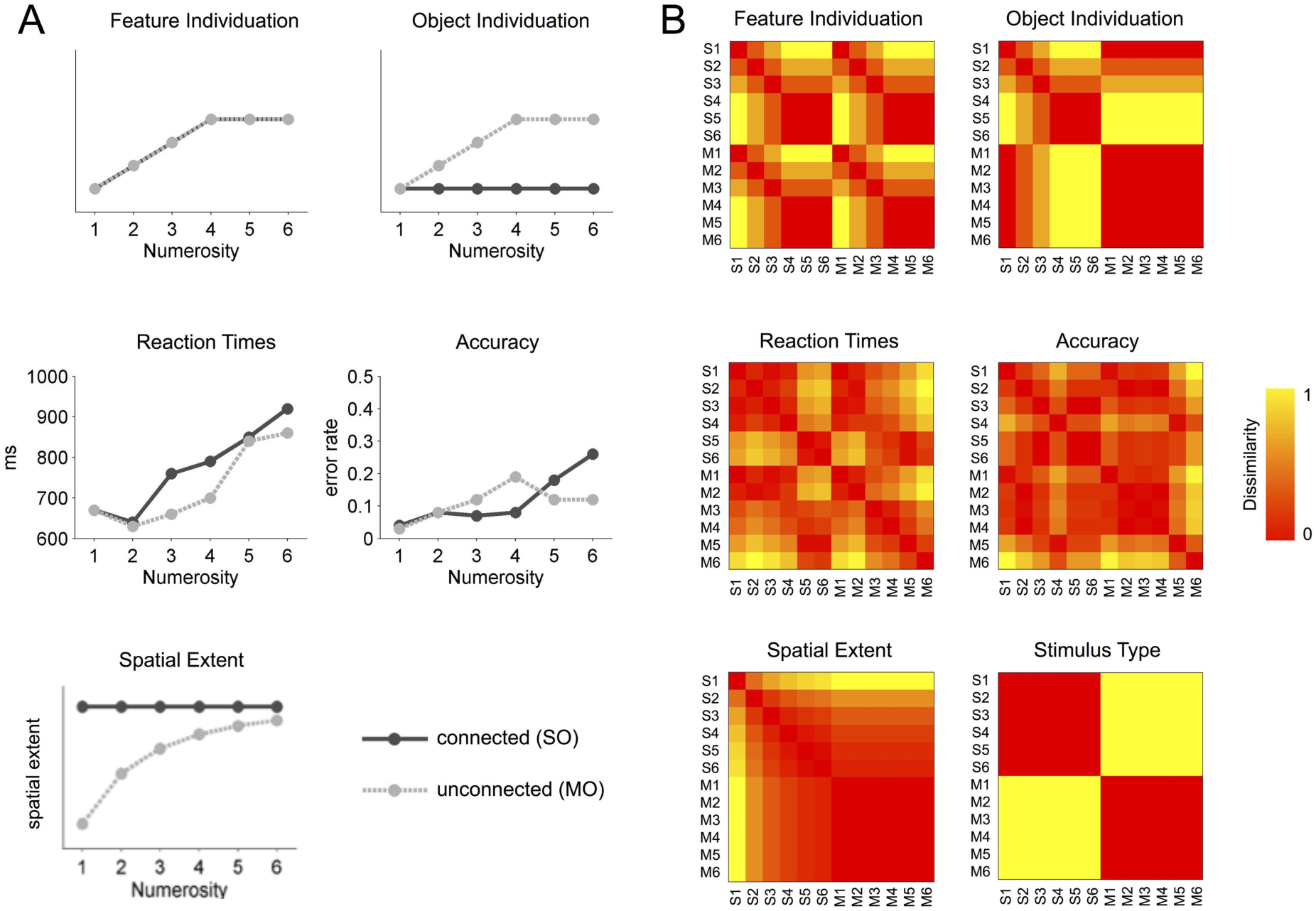
Representational dissimilarity models. (A) Models of neural activity predicted by the feature individuation and object individuation hypotheses (upper row). Mean reaction times and accuracies of the behavioral experiment (lower row; see Methods for details). (B) Representational dissimilarity matrices (RDMs). S1-6: single-object displays (connected features), M1-6: multi-object displays (unconnected features).

The lower triangular parts of the neural and the 6 model RDMs were vectorized, z-scored, and entered as independent and dependent variables, respectively, into a multiple regression. We detected putative collinearity of the models by computing variance inflation factors (VIFs) of the regression coefficients (Belsley, Kuh, & Welsch, 1980). VIFs were < 1.72 suggesting no problematic collinearity between the models. Correlation coefficients were Fisher transformed and entered into one-sample *t*-tests.

For ROI RSA, to identify significant betas, we corrected statistical results (one-sided one sample t tests) for the number of ROIs and tests/models (=18) using the FDR at q = 0.05.

### 2.10 Behavioral Experiment

To compute models of behavioral performance, we collected data on a behavioral discrimination task similar to the task performed in the scanner. This separate behavioral experiment was conducted (rather than using the behavioral responses from the fMRI experiment) for two reasons: First, there were too few oddball blocks per numerosity (n = 7), which might result in unreliable results. Second, and more importantly, we aimed at modeling task difficulty during the non-oddball blocks, which were used in the fMRI analysis. Since there were no responses during these blocks in the fMRI experiment, we used a modified paradigm, in which also the non-oddball blocks had to be responded to. The behavioral experiment used same stimuli as in the fMRI experiment, with new unique stimulus sets created for each participant. Participants (N = 13) performed a number discrimination task consisting of trials with pairs of stimuli. They received instructions to report as quickly and as accurately as possible whether the second stimulus presented in each trial had the same or different number of shapes / arcs as the first stimulus presented. Responses were recorded by button press, with one button representing ‘same’ responses, and another ‘different’.

Subjects completed a short practice round followed by the full-length experiment consisting of 468 trials (pairs of 6 numerosities x 2 stimulus types with 6 presentations per pair). When a pair consisted of different stimulus types, one multi-object and one single-object, the presentation order was counterbalanced with each stimulus taking the first presentation slot 3 times. The order of presentation was randomized for each subject. Each trial consisted of: 1 second fixation, display first stimulus of pair for 500 ms, 600 ms blank gray screen, presentation of second stimulus for 500 ms or until a response was recorded via button press. The presentation timing was designed to mimic the experience of the participants in the fMRI experiment, with the same stimulus duration and inter stimulus interval as the mini-block presentation. Every 40 trials, the participants were given the option to take a self-timed break before continuing.

We recorded reaction times (RT) and accuracy for each trial. For each stimulus pair, we calculated the median RT and entered it into a 12×12 matrix (6 numerosities x 2 stimulus types). To obtain RTs associated with the speed to detect whether a numerosity is the same as in the preceding stimulus, we extracted the on-diagonal of the matrix (the “same” responses) and averaged the 12 RTs across subjects. These RTs were assumed to reflect best the speed to decide *not* to press a button in the non-oddball blocks of the main experiment that were used for fMRI analysis. To compute a representational dissimilarity matrix (RDM) that reflects how dissimilar the different stimuli are in terms of RT, we computed the Euclidean distance between each RT. The same procedure was used to compute an RDM of accuracy.

## 3. Results

### 3.1 Behavioral Results

Participants performed an oddball detection task to ensure they remained attentive throughout the course of the experiment. All participants demonstrated adequate performance, with a hit rate of 0.71 to 0.98 averaging across the 6 runs. Reaction times and accuracies of the behavioral experiment (Figure 4A) were similar to those reported in previous behavioral studies (Porter et al., 2016).

### 3.2 Univariate results

Mean beta estimates for each condition and ROI are plotted in Figure 2. Repeated-measures ANOVA with the factor NUMEROSITY and STIMULUS TYPE revealed significant main effects of numerosity in inferior and superior IPS (both F(1,12) > 9.8, all p < 0.0001) but not in LOC F(1,12) = 0.66, p = 0.65). None of the ROIs showed significant main effects of stimulus type (all F(5,60) < 2.9, all p > 0.11) or interactions (all F(5,60) < 1.33, all p > 0.26). Removing the number of pixels from the regression did not affect the pattern of results, and so are not reported.

We also investigated the point at which neural activation for each additional target plateaued in inferior and superior IPS. Comparisons of the difference in activity from 4 targets to 5, and 5 to 6 in both numerosities suggest that a plateau was established by 5 targets for both single-object and multi-object displays in inferior IPS and LOC and for single-object displays stimuli in superior IPS (Table 1), with the exception that in superior IPS for unconnected targets the effect did not survive FDR correction (q = 0.05; adjusted p = 0.005) but was trending toward significance (t(12) = -2.38, p = 0.035). To test the likelihood in the data that a plateau is reached for sets of items with 5 targets vs. a plateau is reached by 6 targets or more, we performed Bayesian model comparisons (Supplementary Table 1). To this end, we tested a model with linearly increasing weights (1,2,3,4,5,6) and models with linearly increasing weights until a set size of 4 targets (1,2,3,4,4,4) or 5 targets (1,2,3,4,5,5). The ratio of Bayes factors between models reveals how much more likely one model is relative to another model. In both inferior and superior IPS and for both single- and multi-object displays, we found strong evidence (BF ratio > 10) that plateaus were reached with set sizes of 5 than with set sizes of 4. We found no conclusive evidence (BF ratio < 3) regarding whether plateaus were reached with set sizes of 5 or 6 targets (with the exception that in inferior IPS for connected targets a Bayes Factor of 4.8 suggested moderate evidence that a plateau was reached with 6 rather than 5 targets). Together, the Bayesian model comparisons suggest that in inferior and superior IPS a plateau is reached with 5 targets or more rather than 4 targets, but evidence whether a plateau is established by 5 or by 6 targets is inconclusive in this dataset.

**Table 1.** Results of paired t-tests investigating the point at which the increase in percent signal change for each additional target plateaued. Asterisks indicate comparisons that survive FDR correction.

|  | LOC |  | inferior IPS |  | superior IPS |  |
| --- | --- | --- | --- | --- | --- | --- |
|  | t(12) | p | t(12) | p | t(12) | p |
| *four vs. five targets* |  |  |  |  |  |  |
| single-object | -0.281 | 0.783 | -5.243 | <0.001* | -3.452 | 0.005* |
| multi-object | 1.848 | 0.089 | -3.484 | 0.005* | -2.382 | 0.035 |
| *five vs. six targets* |  |  |  |  |  |  |
| single-object | 0.235 | 0.818 | -1.199 | 0.254 | -0.914 | 0.379 |
| multi-object | -0.155 | 0.88 | -1.333 | 0.207 | -0.726 | 0.482 |

### 3.3 Classification of numerosity and stimulus type

Multivoxel pattern classifications were performed to investigate representations of stimulus type and number in the patterns of activity in LOC, inferior IPS, and superior IPS. All ROIs showed classification above chance for numerosity, generalizing across stimulus type (Figure 3A). This suggests all three regions can discriminate number, abstracting across the visual differences between single-object and multi-object displays. LOC also showed significant above chance classification of stimulus type, discriminating between single-object and multi-object displays while generalizing across numerosity (Figure 3C). LOC thus seems to contain additional representations sensitive to the perceptual difference of displays with parts of a single object vs. multiple objects. A repeated measures ANOVA with the factors ROI (LOC, inferior IPS, and superior IPS) and DECODING SCHEME (numerosity, stimulus type; note that the different chance levels of 0.1667 and 0.5 were subtracted from decoding accuracies) revealed an interaction, indicating differential strengths of numerosity decoding (highest in inferior IPS) and stimulus type decoding (highest in LOC; F(2,24) = 5.17, p = 0.014). In addition, a main effect was found for DECODING SCHEME, indicating higher decoding accuracies for numerosity decoding (F(1,12) = 5.34, p = 0.039). Paired t-tests did not reveal significant differences in classification strength between the ROIs after FDR correction (all p > 0.047; q = 0.05; adjusted p = 0.0083).

To identify putative additional regions that encode information about numerosity and stimulus type, we performed a searchlight analysis using identical classification procedures as in the ROI analysis. In line with the ROI analysis we found significant above chance classification of numerosity in bilateral IPS, with the strongest effects in right inferior IPS (Figure 3B, Table 2). Additional regions showing significant classification were lateral occipital cortex, inferior parietal cortex, and lateral prefrontal cortex (bilateral premotor cortex and inferior frontal gyri, right superior frontal gyrus). The stimulus type classification searchlight analysis revealed effects in early visual cortex extending ventrally into lateral occipital, inferior temporal, and fusiform cortices (Figure 3D, Table 2).

**Table 2.** Clusters identified in the classification searchlight analysis.

| Region | x | y | z | t | p | Accuracy |
| --- | --- | --- | --- | --- | --- | --- |
| *Numerosity classification (chance = 16.7%)* |  |  |  |  |  |  |
| left iIPS | -30 | -79 | 32 | 9.76 | <0.000001 | 32.9 |
| right iIPS | 33 | -76 | 32 | 14.59 | <0.000001 | 36.9 |
| right sIPS | 30 | -64 | 43 | 9.69 | 0.000001 | 35.6 |
| left LOC | -27 | -91 | -4 | 9.16 | 0.000001 | 31.0 |
| right LOC | 39 | -82 | 2 | 11.85 | <0.000001 | 30.0 |
| left PMC | -60 | -4 | 38 | 8.62 | 0.000002 | 30.2 |
| right PMC | 39 | -7 | 54 | 9.49 | 0.000001 | 30.0 |
| left SMG | -55 | -46 | 32 | 7.22 | 0.000011 | 27.7 |
| right PoCG | 57 | -36 | 47 | 5.69 | 0.0001 | 30.4 |
| left precuneus | -3 | -67 | 47 | 4.86 | 0.000393 | 25.9 |
| right ITG | 65 | -58 | -16 | 6.63 | 0.000024 | 28.5 |
| left pMTG | -66 | -54 | -4 | 6.74 | 0.000021 | 25.1 |
| *Stimulus Type classification (chance = 50%)* |  |  |  |  |  |  |
| left LOC | -21 | -100 | -1 | 8.85 | 0.000001 | 63.7 |
| right LOC | 24 | -100 | -7 | 10.11 | <0.000001 | 61.8 |
| left ITG/FG | -51 | -79 | -16 | 6.94 | 0.000016 | 64.2 |
| right ITG/FG | 48 | -64 | -26 | 7.09 | 0.000013 | 62.3 |

Coordinates (x, y, z) in MNI space. Maps were thresholded at $p = 0.001$ , corrected cluster threshold $p = 0.05$ . Abbreviations: FG, fusiform gyrus; i/pIPS, inferior/posterior intraparietal sulcus; ITG, inferior temporal gyrus; LOC, lateral occipital cortex; SMG, supramarginal gyrus; PMC, premotor cortex; pMTG, posterior middle temporal gyrus; PoCG, postcentral gyrus; SMG, supramarginal gyrus; SPL, superior parietal lobe.

Are there brain areas that are particularly tuned to discriminate numerosities of connected as compared to unconnected stimuli, and vice versa? We addressed this question by decoding numerosity within each stimulus type separately. Both connected and unconnected stimulus types could be decoded in the network of parietal, prefrontal, and occipital areas described above (Supplementary Fig. 4). Contrasting the numerosity decoding maps for multi-object and single-object displays against each other did not reveal significant clusters, suggesting that the neural mechanisms involved in individuating numerosities of parts of a single object and multiple distinct objects do not substantially differ.

### 3.4 Representational similarity analysis

To test the two individuation models under investigation more directly, we performed a representational similarity analysis (RSA) that compared the similarities between the neural patterns associated with the conditions with model-based similarities predicted by the Feature Individuation and Object Individuation hypotheses. Specifically, the Feature Individuation model assumes that numerosities are represented similarly for multi- and single-object displays (with distinguishable activation patterns in the subitizing range from 1-4 and indistinguishable patterns beyond the subitizing range), whereas the Object Individuation model assumes that single-object displays are represented similarly across numerosities and similar to the multi-object display with numerosity = 1 (because all show one separable object) and only the single-object displays show distinguishable activation patterns in the subitizing range from 1-4 and indistinguishable patterns beyond the subitizing range (Figure 4). In each of the ROIs we correlated the neural activity patterns with each other resulting in a 12 x 12 correlation matrix that was converted into a neural representational dissimilarity matrix (RDM; 1 – r). The neural RDM was entered into a multiple regression with predictors for Feature Individuation, Object Individuation, Spatial Extent, Stimulus Type, and two behavioral models - Reaction Times and Accuracy – that were based on a separate behavioral experiment (Figure 4A; see *Methods, Behavioral Experiment*, for details). The Spatial Extent model was included to capture perceptual and attentional effects associated with the spatial extent of the area in which the targets could occur. Note that the Spatial Extent model resembles to some extent the Stimulus Type model, which, however, captures a categorical distinction between multi- and single-object displays. The Reaction Times and Accuracy models were included to test whether the similarity of neural activity patterns could be explained by similarity of conditions in terms of task difficulty.

In all ROIs, variance in neural dissimilarity could be explained by the Feature Individuation model independently of other tested models (Figure 5A). In all ROIs, additional variance could be explained by the Reaction Times model. In LOC and inferior IPS, additional variance was explained by the Spatial Extent model. In none of the ROIs did the Object Individuation, Stimulus Type, and Accuracy models show significant effects (following FDR correction; q = 0.05, adjusted p = 0.015). Paired t-tests revealed that in all ROIs the Feature Individuation model could explain significantly more variance than the Object Individuation model (all t(12) > 4.71, p < 0.001). Paired t-tests between ROIs revealed that the Spatial Extent model explained significantly more variance in LOC vs. superior IPS (t(12) > 3.68, p = 0.003).

**Figure 5.**
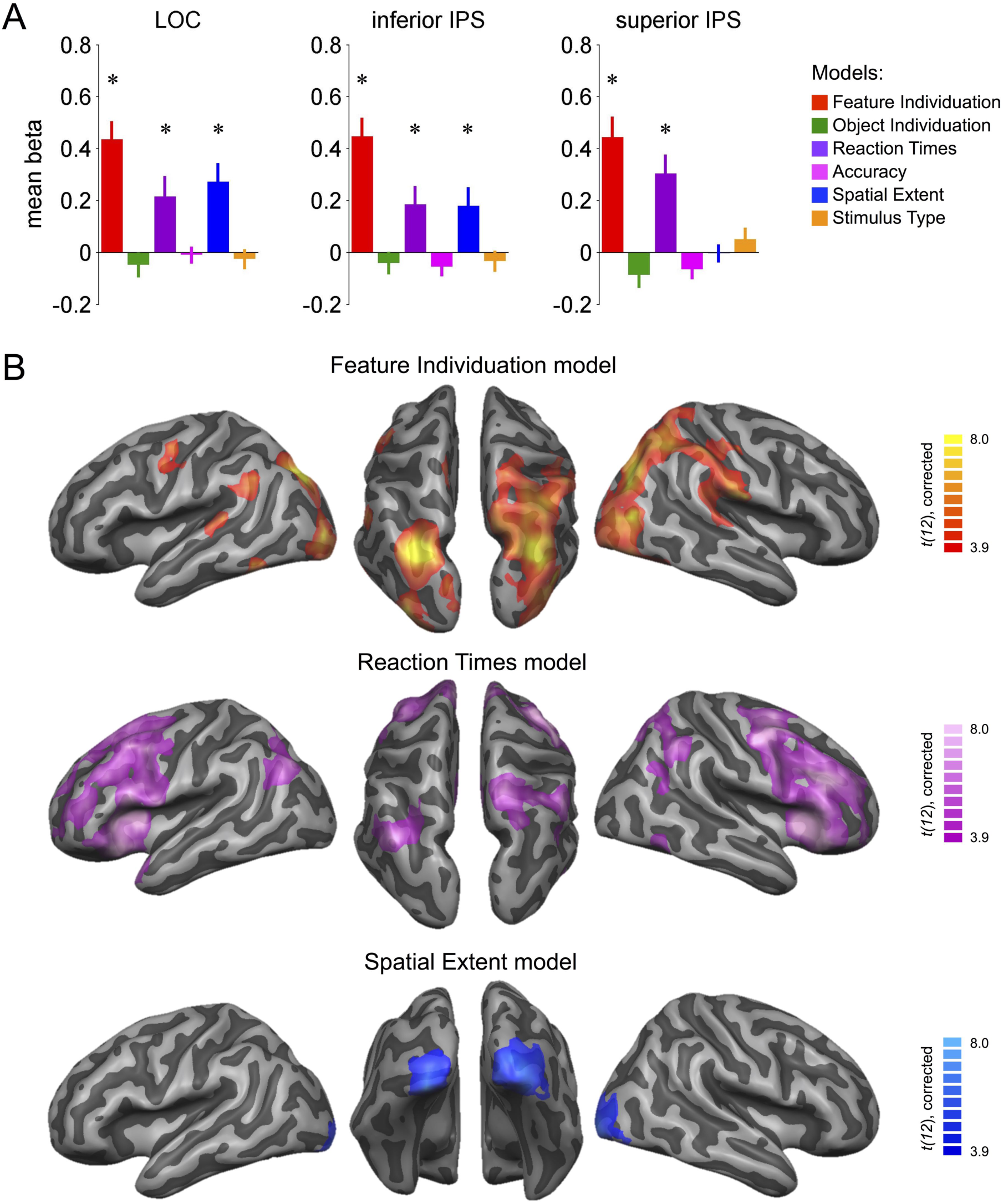
Multiple regression RSA. (A) ROI analysis. Bars show parameter estimates of the multiple regression analysis relating model RDMs to neural RDMs of the ROIs. Asterisks indicate FDR corrected significant beta estimates; error bars indicate SEM. (B) Statistical maps for each model of the searchlight multiple regression RSA (cluster size corrected for multiple comparisons at p = 0.05, initial voxel threshold p = 0.001). Maps of the Object Individuation and Accuracy models did not reveal clusters that survived correction for multiple comparisons.

To identify other areas in which representational organization can be explained by the Feature and Object Individuation models, we performed a searchlight multiple regression RSA (Figure 5B, Table 3). The Feature Individuation model revealed clusters in bilateral IPS, LOC, and supramarginal gyrus, left premotor cortex, left postcentral sulcus, left superior temporal gyrus, right superior IPS, and right inferior temporal cortex. Effects were strongest in a distinct peak area in inferior IPS bilaterally. Notably, this area was close to, but not perfectly overlapping with the inferior IPS ROI (Supplementary Figure 1A), which might explain why the ROI analysis did not reveal significantly stronger effects for the Feature Individuation model in inferior as compared to superior IPS and LOC. The Spatial Extent model revealed effects in early visual cortex. Additional effects were found in right inferior IPS, which, however, did not survive correction for multiple comparisons (p < 0.005, uncorrected). The Reaction Times model revealed the strongest effects in bilateral frontal operculum, inferior frontal gyrus, and insula as well as in dorsolateral prefrontal cortex and superior IPS/angular gyrus. The Object Individuation, Stimulus Type, and Accuracy models did not reveal significant clusters, even after applying more liberal thresholds. However, a trend was observed for the Stimulus Type model in bilateral inferior temporal gyrus (p < 0.001, uncorrected). The absence of effects for the Stimulus Type model in early visual cortex does not contradict the results of the Stimulus Type decoding, because lower-level perceptual differences between the two stimulus displays were better captured by the Spatial Extent model.

**Table 3.** Clusters identified in the multiple regression searchlight RSA that included models of Feature Individuation, Object Individuation, Stimulus Type, Reaction Times, and Accuracy.

| Region | x | y | z | t | p |
| --- | --- | --- | --- | --- | --- |
| *Feature Individuation* |  |  |  |  |  |
| left iIPS | -27 | -76 | 32 | 13.53 | <1.00E-06 |
| right iIPS | 30 | -70 | 26 | 10.31 | <1.00E-06 |
| right sIPS | 39 | -55 | 50 | 6.84 | 1.80E-05 |
| left LOC | -42 | -91 | -4 | 8.11 | 3.00E-06 |
| right LOC | 45 | -76 | 8 | 8.30 | 3.00E-06 |
| left PMC | -45 | -7 | 50 | 5.76 | 9.00E-05 |
| left SMG | -51 | -46 | 26 | 6.12 | 5.20E-05 |
| right SMG | 63 | -22 | 26 | 6.63 | 2.40E-05 |
| right sIPS | 39 | -55 | 50 | 6.84 | 1.80E-05 |
| left STG | -57 | -34 | 11 | 6.27 | 4.10E-05 |
| left PoCS | -21 | -34 | 62 | 8.97 | 1.00E-06 |
| right ITG | -57 | -64 | -22 | 6.99 | 1.50E-05 |
| *Spatial Extent* |  |  |  |  |  |
| left EVC | -21 | -94 | -16 | 7.64 | 6.00E-06 |
| right EVC | 21 | -85 | -19 | 8.73 | 1.00E-06 |
| *Reaction Times* |  |  |  |  |  |
| left FOP/IFG | -39 | 8 | 5 | 11.12 | <1.00E-06 |
| right FOP/IFG | 36 | 26 | -1 | 15.92 | <1.00E-06 |
| left DLPFC | -42 | 14 | 44 | 8.43 | 2.00E-06 |
| right DLPFC | 45 | 26 | 26 | 12.27 | <1.00E-06 |
| left SMA | -6 | 20 | 47 | 10.63 | <1.00E-06 |
| right SMA | 9 | 17 | 44 | 15.22 | <1.00E-06 |
| left AG | -39 | -76 | 41 | 7.30 | 9.00E-06 |
| right AG | 36 | -55 | 47 | 6.98 | 1.50E-05 |
| right SPL | 21 | -67 | 66 | 9.37 | 1.00E-06 |
| right PMC | 36 | 5 | 53 | 9.91 | <1.00E-06 |
| right ITG | 51 | -64 | -19 | 6.78 | 2.00E-05 |
| lingual gyrus | 15 | -64 | 2 | 9.42 | 1.00E-06 |

Coordinates (x, y, z) in MNI space. Maps were thresholded at $p = 0.001$ , corrected cluster threshold $p = 0.05$ . Abbreviations: AG, angular gyrus; DLPFC, dorsolateral prefrontal cortex; EVC, early visual cortex; FOP, frontal operculum; IFG, inferior frontal gyrus; i/sIPS, inferior/superior intraparietal sulcus; ITG, inferior temporal gyrus; LOC, lateral occipital cortex; SMG, supramarginal gyrus; PMC, premotor cortex; PoCS, postcentral sulcus; SMA, supplementary motor area; SMG, supramarginal gyrus; SPL, superior parietal lobe; STG, superior temporal gyrus.

To further inspect the representational organization along numerosity and stimulus type in the tested ROIs, we performed a cluster analysis (Supplementary Fig. 5). This analysis revealed that numerosities clustered together independently of stimulus type in all ROIs, but more strongly in superior IPs and less strongly in LOC. This result provides another demonstration that neural representations of numerosity are similar irrespective of connectedness.

## 4. Discussion

In this experiment, we aimed to test whether individuation of connected object parts and unconnected objects rely on a common mechanism or independent neural mechanisms. We report three main findings. Firstly, we show that the number of targets, regardless of connectedness, modulates neural activity in IPS. This result corroborates previous findings of EEG modulation in response to individuation of connected parts of single objects (Poncet et al., 2016) and extends them by providing a direct comparison of neural response patterns for individuation of parts of a single object and multiple distinct objects. Secondly, and most importantly, multi-voxel pattern analyses investigating the representation of information within these regions demonstrate that inferior and superior IPS as well as LOC hold representations of numerosity that generalize across stimulus type. By contrast, only the LOC performed above chance when discriminating multi-object and single-object displays. Thirdly, the neural representation of the tested stimulus items was best explained by a model of Feature Individuation that does not assume any difference in individuation between the two stimulus types. Moreover, the representation of numerosity was independent from effects of task difficulty. Searchlight analyses aimed at detecting effects of numerosity and stimulus type in the whole brain corroborated the results of the ROI analysis but revealed more distinctive effects by demonstrating that numerosity effects were strongest in a well-circumscribed region in inferior IPS in both hemispheres. Together, these findings provide clear support for the hypothesis that parallel individuation of object parts and distinct objects relies on a common neural mechanism, which draws on the inferior IPS. Objecthood is therefore not a necessary condition to detect small numbers at once; inferior IPS individuates the parts of a single object with the same ease and neural signatures as multiple separate objects.

Multivoxel pattern analyses allowed us to look at whether neural response patterns in our regions of interest as well as in other cortical regions held information about numerosity or stimulus type, with the ability to generalize across the other dimension. Both ROI and searchlight analyses revealed that inferior and superior IPS as well as LOC contain representations that discriminate numerosities, generalizing across multi-object and single-object stimuli. By contrast, only LOC could classify the two stimulus types independently of numerosity. The result of stimulus-general decoding of numerosity provides an important extension of previous findings, with one study showing that voxels in the parietal lobe could classify number generalizing across changes in spatial configurations, as well as modality (Damarla et al., 2016), and another demonstrating that the posterior parietal cortex can classify number in both enumeration and visual working memory tasks (Knops et al., 2014). Additionally, the inferior IPS has been shown to be able to decode both the number of objects in a display as well as discriminate between displays either consisting of multiple objects of the same shape, or unique shapes (Naughtin, Mattingley, & Dux, 2016). Our findings demonstrate that IPS represents numerosity irrespective of connectedness, which suggests a common neural mechanism for parallel individuation of both connected parts and unconnected objects in IPS. By contrast, our findings argue against the hypothesis that parallel individuation in IPS applies only to single, unconnected objects, as suggested by previous studies (He et al., 2015; Scholl et al., 2001; Xu & Chun, 2007). Moreover, we found no evidence for brain areas that specifically represent numerosity of either object parts or independent objects. Notably, we also observed that LOC discriminated numerosities irrespective of the specific stimulus display but did not show effects of numerosity on univariate neural activation typically associated with individuation in the subitizing range. We therefore conclude that LOC is not primarily involved in object individuation but rather discriminated numerosities based on perceptual details that were common across connected and unconnected stimulus displays, i.e., the arcs that varied with numerosity in both stimulus types in a similar way. This interpretation is in line with the finding that visual cortex responds to numerosity of isolated and connected dots in a similar manner, which suggests that visual areas represent perceptual aspects of numerosity prior to later segmentation processes (Fornaciai & Park, 2018). Thus, together with the finding that LOC discriminated between the two stimulus displays in general, it appears that LOC is sensitive to perceptual aspects of numerosity whereas higher level representations of numerosity are captured by IPS only. Notably, effects of numerosity in LOC – and even in more posterior visual areas as revealed by the searchlight analyses – cannot be explained by low-level visual effects because the exact locations of the targets varied across stimulus exemplars. This finding is in line with the demonstration that early visual cortex is sensitive to numerosity independently of lower-level aspects like extent, density, size, and location of targets (DeWind et al., 2019). However, numerosities do not appear to be decodable across visual and auditory modalities in visual cortex (Cavdaroglu et al., 2015), suggesting that numerosity representation in ventral areas is tied to the visual modality.

To explore what kind of information is represented in the ROIs with higher precision, we compared the neural representational similarity of the voxel patterns associated with each stimulus to several different models. Notably, this approach allowed us to test hypothesized individuation models independently of task difficulty. ROI and searchlight analyses consistently found that the Feature Individuation model – which postulates that multiple item individuation can occur both over objects and features of an object – explained the neural variance in inferior and superior IPS and LOC significantly better than the (separate) Object Individuation model, which postulates that objecthood is constitutive for individuation and therefore an object with connected features is treated as a single object and doesn’t allow multiple feature individuation. Importantly, while additional variance in neural representation could be explained by task difficulty (as modeled by reaction time in a behavioral numerosity discrimination task), the effects of the Feature Individuation model were independent of the effect of task difficulty. This finding is in line with the observation that activity in the inferior IPS is modulated by the number of targets even when conditions were matched for task difficulty (Cusack, Mitchell, & Duncan, 2010). In LOC and inferior IPS representational similarity could be explained by a model of spatial extent of the stimuli. This model was intended to capture lower-level visual effects and possibly effects related to spatial attention. The searchlight analysis revealed that effects of spatial extent were strongest in early visual cortex and extended only into lateral occipital and posterior parietal areas known to contain retinotopic organization (Sereno et al., 1995; Swisher, Halko, Merabet, McMains, & Somers, 2007). Notably, there was no distinct cluster in inferior IPS but rather the effect gradually spread into IPS. Most importantly, effects of feature individuation were over and above effects related to spatial extent of the stimuli. Spatial extent alone is therefore not sufficient to explain effects of numerosity in inferior IPS. Notably, while effects of spatial extent peaked in areas posterior of inferior IPS, effects of task difficulty peaked more anteriorly and superiorly in parietal cortex. Together, these findings point toward a posterior-anterior gradient of functions along IPS: Stimulus-related information as captured by the spatial extent model is represented in posterior areas such as LOC. Information processed in LOC serves as input to inferior IPS, which captures effects related to feature/object, individuation/numerosity representations. Finally, more anterior areas in IPS as well as in prefrontal cortex show effects of task difficulty and thus appear to encode information at the output level. Following this interpretation, the question arises whether numerosity representation in dorsal areas, specifically in inferior and/or superior IPS, depends on explicit attention toward numerosity of specific targets. While visual areas have been shown to represent numerosity in a task-independent manner, numerosity could not be decoded in parietal areas when numerosity was irrelevant for the task (Cavdaroglu et al., 2015; Fornaciai & Park, 2018). It remains to be tested whether IPS would represent single-object displays as a single object (or represent numerosity at all) if this information was irrelevant for the task. In particular for ambiguous scenarios – which is usually the default in natural scenes that contain different object ensembles and objects, which may have different parts and subparts – it appears plausible that numerosity representation depends on attention toward specific targets. Consequently, such flexible representation of numerosity must be independent of the specific stimulus. Indeed, behavioral adaptation studies demonstrated the existence of stimulus-independent numerosity representations that generalize across format (e.g. dot arrays vs. sequences of stimuli) or modality (visual vs. auditory) (Arrighi, Togoli, & Burr, 2014) and neuroimaging studies provided evidence for stimulus-independent number representations in IPS (Damarla et al., 2016; Piazza, Pinel, Le Bihan, & Dehaene, 2007). Our finding that decoding of numerosity across single- and multi-object displays was strongest in inferior IPS supports the view that this area represents numerosity in a flexible (= task-dependent) and stimulus-independent manner. Since our task required attention toward the number of targets in single- and multi-object displays, it might not come as a surprise that we found a common “abstract” representation of numerosity in IPS. The goal of the study, however, was to test whether individuation of connected object parts and unconnected objects relies on independent mechanisms or not. In our study we cannot dissociate object individuation from numerosity representation. However, numerosity decoding across displays and the absence of differences between numerosity decoding of single- and multi-object displays throughout the pathway from visual areas to IPS suggests that not only numerosity representation but also the individuation mechanism was similar for the two displays. From a more general perspective, it appears plausible that this individuation mechanism is functionally related to attentional mechanisms (Corbetta & Shulman, 2002) as well as to numerical cognition (Nieder & Dehaene, 2009), all of which associated with IPS.

Previous studies have suggested that grouping and connectedness can affect the neural individuation and enumeration of items (He et al., 2015; Xu & Chun, 2007). These studies demonstrated main effects of grouping and connectedness in inferior IPS, which is in contrast to our finding of similar activation profiles for single- and multi-object displays in IPS. Xu & Chun (2007) reported that when shapes were grouped by a common background, they evoked smaller response magnitudes than a similar display with each shape occurring on its own background. Another study showed that connecting a subset of dots in a display caused underestimation in behavioral judgments, as well as a shift in the neural adaptation curves in IPS indicative of underestimation (He et al., 2015). Based on these results, one might expect the response of each numerosity in the single-object displays with connected features to be lower than that of the comparable multi-object display. The lack of this univariate effect in the current study could be the result of different selection demands. Grouping in Xu & Chun’s (2007) study was performed by placing objects within one of several dark fields present in the gray display, thus even when all targets were grouped within the same field, other black fields were still visible against the gray background. The selection of figure could shift between the larger object, the dark field, or the smaller feature, the target shape. This was also true of the displays used in He et al. (2015). Their task required participants to make judgments about the number of dots in a display. These displays also included irrelevant lines, which could sometimes be oriented so that they connected some of the dots. In their connected condition, the same display would thus contain two possible definitions of figure and ground, both of which were necessary to accurately perform the task. We suggest that the effects in IPS are not a direct result of connectedness, but rather the addition of a second and confounding figure/ground factor within the same display: In displays showing dots of which some are connected by lines, dots may be represented as figure (and the lines are perceived as ground), or vice versa. If the task is to enumerate the dots, then the lines are perceived as ground that may spatially overlap with the dots, which might impede individuation of the dots. This interpretation would also explain the behavioral finding that numerosities of connected dots are systematically underestimated (Fornaciai et al., 2016; Franconeri et al., 2009; He et al., 2009). By contrast, our single-object stimuli only contained one relevant definition of figure and ground to perform the task – the protruding arcs were the target figures against a circular ground. Therefore, the decrease in activity previously observed in the inferior IPS to connected and grouped items may have resulted from competing levels of selection within the same task, with a bias toward figure as defined by a lack of connectedness. In contrast, when connectedness had a uniform effect on the selection of targets, as in our study, connected features could be individuated independently and in parallel. Thus, our results are generally consistent with the neural object file theory proposed in Xu & Chun (2009), suggesting that the inferior IPS contributes to feature/object individuation. We propose to add to the neural object file theory the specification that the inferior IPS can operate over a flexible definition of figure and ground. When figure and ground are not confounded, connectedness does not have a detrimental effect on multiple feature individuation performance.

In conclusion, we observed similar neural signatures of parallel individuation for both unconnected, separate objects and connected features of a single object. The neural similarity of numerosity representations in IPS suggest that individuation of parts of a single object and multiple objects relies on a common neural mechanism that is distinct from visual processing of objects and object parts in LOC.

## 5. Acknowledgements

This work was supported by the Fondazione CARITRO and the Provincia Autonoma di Trento to AC. We thank Valentina Brentari, Wilhelm Malloni, and Mark Greenlee for their help in data collection and Yaoda Xu and Veronica Mazza for their advice on this project.

## 7. Data and Code Availability and Pre-registration Statement

Neuroimaging data, stimulus material, and code are available at https://osf.io/56yz7/.

## Footnotes

1 Note that here we use the term connectedness instead of connectivity to avoid terminological confusion with the concept of neural connectivity in the neuroimaging literature.

## References

Akin, O., & Chase, W. (1978). Quantification of three-dimensional structures. J Exp Psychol Hum Percept Perform, 4(3), 397–410.

Ansari, D., Lyons, I. M., van Eimeren, L., & Xu, F. (2007). Linking visual attention and number processing in the brain: the role of the temporo-parietal junction in small and large symbolic and nonsymbolic number comparison. J Cogn Neurosci, 19(11), 1845–1853.

Arrighi, R., Togoli, I., & Burr, D. C. (2014). A generalized sense of number. Proc Biol Sci, 281(1797).

Belsley, D. A., Kuh, E., & Welsch, R. E. (1980). Regression diagnostics: Identifying influential data and sources of collinearity (Vol. 571): John Wiley & Sons.

Cavdaroglu, S., Katz, C., & Knops, A. (2015). Dissociating estimation from comparison and response eliminates parietal involvement in sequential numerosity perception. Neuroimage, 116, 135–148.

Corbetta, M., & Shulman, G. L. (2002). Control of goal-directed and stimulus-driven attention in the brain. Nat Rev Neurosci, 3(3), 201–215.

Cusack, R., Mitchell, D. J., & Duncan, J. (2010). Discrete object representation, attention switching, and task difficulty in the parietal lobe. J Cogn Neurosci, 22(1), 32–47.

Cutini, S., Scatturin, P., Basso Moro, S., & Zorzi, M. (2014). Are the neural correlates of subitizing and estimation dissociable? An fNIRS investigation. Neuroimage, 85 Pt 1, 391–399.

Damarla, S. R., Cherkassky, V. L., & Just, M. A. (2016). Modality-independent representations of small quantities based on brain activation patterns. Hum Brain Mapp, 37(4), 1296–1307.

DeWind, N. K., Park, J., Woldorff, M. G., & Brannon, E. M. (2019). Numerical encoding in early visual cortex. Cortex, 114, 76–89.

Drew, T., & Vogel, E. K. (2008). Neural measures of individual differences in selecting and tracking multiple moving objects. J Neurosci, 28(16), 4183–4191.

Driver, J., & Vogel, E. K. (1998). Attention and visual object segmentation. In R. Parasuraman (Ed.), The attentive brain (pp. 299–325). Cambridge, MA: MIT Press.

Ester, E. F., Drew, T., Klee, D., Vogel, E. K., & Awh, E. (2012). Neural measures reveal a fixed item limit in subitizing. J Neurosci, 32(21), 7169–7177.

Fornaciai, M., Cicchini, G. M., & Burr, D. C. (2016). Adaptation to number operates on perceived rather than physical numerosity. Cognition, 151, 63–67.

Fornaciai, M., & Park, J. (2018). Early Numerosity Encoding in Visual Cortex Is Not Sufficient for the Representation of Numerical Magnitude. J Cogn Neurosci, 1–15.

Franconeri, S. L., Bemis, D. K., & Alvarez, G. A. (2009). Number estimation relies on a set of segmented objects. Cognition, 113(1), 1–13.

Friston, K. J., Fletcher, P., Josephs, O., Holmes, A., Rugg, M. D., & Turner, R. (1998). Event-related fMRI: characterizing differential responses. Neuroimage, 7(1), 30–40.

He, L. X., Zhang, J., Zhou, T., & Chen, L. (2009). Connectedness affects dot numerosity judgment: implications for configural processing. Psychon Bull Rev, 16(3), 509–517.

He, L. X., Zhou, K., Zhou, T. G., He, S., & Chen, L. (2015). Topology-defined units in numerosity perception. Proceedings of the National Academy of Sciences of the United States of America, 112(41), E5647–E5655.

Howe, P. D. L., Cohen, M. A., Pinto, Y., & Horowitz, T. S. (2010). Distinguishing between parallel and serial accounts of multiple object tracking. Journal of Vision, 10(8).

Hubbard, E. M., Piazza, M., Pinel, P., & Dehaene, S. (2005). Interactions between number and space in parietal cortex. Nature Reviews Neuroscience, 6(6), 435–448.

Humphreys, G. W., & Riddoch, M. J. (1993). Interactions between Object and Space Systems Revealed through Neuropsychology. Attention and Performance, 14, 143–162.

Jeffreys, H. (1961). Theory of probability, (Oxford: Oxford University Press).

Kahneman, D., Treisman, A., & Gibbs, B. J. (1992). The reviewing of object files: object-specific integration of information. Cogn Psychol, 24(2), 175–219.

Kaufman, E. L., Lord, M. W., Reese, T. W., & Volkmann, J. (1949). The discrimination of visual number. Am J Psychol, 62(4), 498–525.

Knops, A., Piazza, M., Sengupta, R., Eger, E., & Melcher, D. (2014). A Shared, Flexible Neural Map Architecture Reflects Capacity Limits in Both Visual Short-Term Memory and Enumeration. Journal of Neuroscience, 34(30), 9857–9866.

Leslie, A. M., Xu, F., Tremoulet, P. D., & Scholl, B. J. (1998). Indexing and the object concept: developing ‘what’ and ‘where’ systems. Trends Cogn Sci, 2(1), 10–18.

Mazza, V., & Caramazza, A. (2011). Temporal brain dynamics of multiple object processing: the flexibility of individuation. PLoS One, 6(2), e17453.

Mazza, V., & Caramazza, A. (2015). Multiple object individuation and subitizing in enumeration: a view from electrophysiology. Front Hum Neurosci, 9, 162.

Mazza, V., Pagano, S., & Caramazza, A. (2013). Multiple object individuation and exact enumeration. J Cogn Neurosci, 25(5), 697–705.

Mishkin, M., Ungerleider, L. G., & Macko, K. A. (1983). Object Vision and Spatial Vision - 2 Cortical Pathways. Trends in Neurosciences, 6(10), 414–417.

Mitchell, D. J., & Cusack, R. (2008). Flexible, capacity-limited activity of posterior parietal cortex in perceptual as well as visual short-term memory tasks. Cerebral Cortex, 18(8), 1788–1798.

Morey, R. D., Rouder, J. N., Jamil, T., & Morey, M. R. D. (2015). Package ‘BayesFactor’. URL http://cran.r-project.org/web/packages/BayesFactor/BayesFactor.pdf (accessed 10.06. 15).

Naughtin, C. K., Mattingley, J. B., & Dux, P. E. (2016). Distributed and Overlapping Neural Substrates for Object Individuation and Identification in Visual Short-Term Memory. Cerebral Cortex, 26(2), 566–575.

Nieder, A., & Dehaene, S. (2009). Representation of number in the brain. Annu Rev Neurosci, 32, 185–208.

Oosterhof, N. N., Connolly, A. C., & Haxby, J. V. (2016). CoSMoMVPA: Multi-Modal Multivariate Pattern Analysis of Neuroimaging Data in Matlab/GNU Octave. Front Neuroinform, 10, 27.

Pagano, S., Lombardi, L., & Mazza, V. (2014). Brain dynamics of attention and working memory engagement in subitizing. Brain Res, 1543, 244–252.

Pagano, S., & Mazza, V. (2012). Individuation of multiple targets during visual enumeration: new insights from electrophysiology. Neuropsychologia, 50(5), 754–761.

Palmer, S., & Rock, I. (1994). Rethinking perceptual organization: The role of uniform connectedness. Psychon Bull Rev, 1(1), 29–55.

Piazza, M., Pinel, P., Le Bihan, D., & Dehaene, S. (2007). A magnitude code common to numerosities and number symbols in human intraparietal cortex. Neuron, 53(2), 293–305.

Poncet, M., Caramazza, A., & Mazza, V. (2016). Individuation of objects and object parts rely on the same neuronal mechanism. Sci Rep, 6, 38434.

Porter, K. B., Mazza, V., Garofalo, A., & Caramazza, A. (2016). Visual object individuation occurs over object wholes, parts, and even holes. Attention Perception & Psychophysics, 78(4), 1145–1162.

Pylyshyn, Z. W. (1989). The role of location indexes in spatial perception: a sketch of the FINST spatial-index model. Cognition, 32(1), 65–97.

Pylyshyn, Z. W., & Storm, R. W. (1988). Tracking multiple independent targets: evidence for a parallel tracking mechanism. Spat Vis, 3(3), 179–197.

Sathian, K., Simon, T. J., Peterson, S., Patel, G. A., Hoffman, J. M., & Grafton, S. T. (1999). Neural evidence linking visual object enumeration and attention. Journal of Cognitive Neuroscience, 11(1), 36–51.

Scholl, B. J., Pylyshyn, Z. W., & Feldman, J. (2001). What is a visual object? Evidence from target merging in multiple object tracking. Cognition, 80(1-2), 159–177.

Sereno, M. I., Dale, A. M., Reppas, J. B., Kwong, K. K., Belliveau, J. W., Brady, T. J., … Tootell, R. B. (1995). Borders of multiple visual areas in humans revealed by functional magnetic resonance imaging. Science, 268(5212), 889–893.

Swisher, J. D., Halko, M. A., Merabet, L. B., McMains, S. A., & Somers, D. C. (2007). Visual topography of human intraparietal sulcus. J Neurosci, 27(20), 5326–5337.

Todd, J. J., & Marois, R. (2004). Capacity limit of visual short-term memory in human posterior parietal cortex. Nature, 428(6984), 751–754.

Trick, L. M., & Pylyshyn, Z. W. (1994). Why Are Small and Large Numbers Enumerated Differently - a Limited-Capacity Preattentive Stage in Vision. Psychol Rev, 101(1), 80–102.

Xu, Y. (2009). Distinctive neural mechanisms supporting visual object individuation and identification. J Cogn Neurosci, 21(3), 511–518.

Xu, Y., & Chun, M. M. (2006). Dissociable neural mechanisms supporting visual short-term memory for objects. Nature, 440(7080), 91–95.

Xu, Y., & Chun, M. M. (2007). Visual grouping in human parietal cortex. Proc Natl Acad Sci U S A, 104(47), 18766–18771.

Xu, Y., & Chun, M. M. (2009). Selecting and perceiving multiple visual objects. Trends Cogn Sci, 13(4), 167–174.

