## Supplementary Materials for "Individuation of parts of a single object and multiple distinct objects relies on a common neural mechanism in inferior intraparietal sulcus"

### Supplementary Material

#### Supplementary Methods

*Definition of individually defined ROIs*

To define individual ROIs in the participants, two localizer scans were acquired. To define the LOC and inferior IPS, we used an object localizer as used in Xu & Chun (2006) and Xu (2009). This localizer consisted of blocks of displays consisting of six black shapes on a gray background, and then blocks of noise images. Each image was presented for 500 ms with an inter-stimulus-interval (ISI) of 200 ms. Subjects performed a motion detection task. Each subject performed 2 runs, each 4.67 minutes long. We decided to not modify the shapes used in this localizer to reflect the stimuli used in our main experiment since there were not enough black pixels in the outline shape to generate noise images. To define superior IPS, we used the visual working memory (VSTM) task from Xu (2009) with the exclusion of the identical shape trials; all shapes presented in each display were unique. One, 2, 3, 4, or 6 black shapes were presented around fixation for 200 ms, creating a display very similar to those used as the shape stimuli from the first localizer, followed by a 1 s blank, after which a test shape was presented at fixation, and the subject pressed a button to indicate whether the shape had been present in the display (index finger), or absent from the previous display (middle finger). Subjects had 2.5s to respond before receiving feedback for that trial. A feedback display was shown for 1.3 s, indicating whether the subject's response was correct (smiley face presented at fixation) or incorrect (sad face). Each subject performed 2 runs of this task, each lasting 7.97 minutes. We also decided not to use our outline stimuli as the shapes in the VSTM task, since they are so visually similar we expected it would be too difficult for subjects to perform.

To localize the inferior IPS and LOC, we performed a general linear model with two regressors: objects and noise. The two ROIs were defined based on a contrast of objects > noise. The superior IPS was localized based on a general linear model with each set size of the VSTM task weighted by the Cowan's K estimate for that set size for that participant. The K-value was calculated as $K = (HR+CR-1)*N$ where $K$ = number of items encoded, HR = hit rate, CR = correct rejection rate, and N = set size. All three regions of interest were defined individually for each participant. For each region, the peak voxel with the highest beta value was located within the cluster closest to the peak coordinates as previously reported for these regions (Todd and Marois 2004; Xu 2009). For univariate analyses, a 5.5 mm radius used; for multivariate analyses, a ROI size of 9 mm radius sphere was used. ROIs were averaged across hemisphere. LOC and inferior IPS could be identified in all 13 participants. The left superior IPS could be identified in 12 of 13 participants, the right superior IPS could be identified in 11 of 13 participants.

Mean MNI coordinates (x, y, z, standard deviations in parentheses; see also Supplementary Figure 1A) of the ROIs were as follows: left LOC: -30 (5.4), -83 (5.7), 20 (2.5); right LOC: 37 (10.1), -77 (9.4), 18 (8.8); left inferior IPS: -26 (8.0), -75 (11.2), 36 (7.0); right inferior IPS: 27 (5.3), -73 (9.0), 38 (5.4); left superior IPS: -26 (6.5), -62 (4.8), 47 (7.3); right superior IPS: 31 (9.4), -59 (5.2), 51 (6.8).

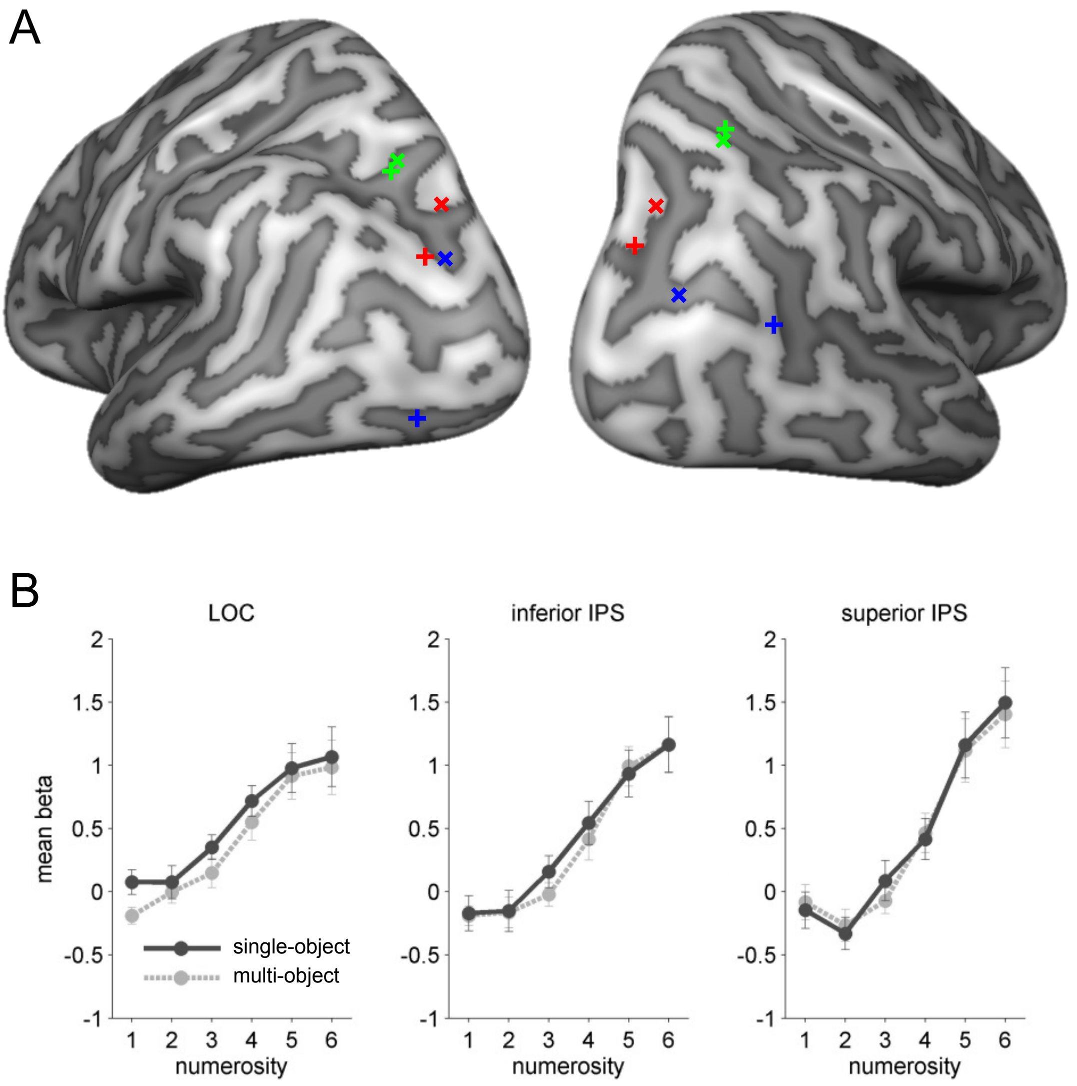


**Supplementary Figure 1.** (A) Comparison between group-averaged individually defined ROIs (indicated by x) and ROIs based on Xu (2009; indicated by +). Green: superior IPS, red: inferior IPS, blue: LOC. (B) Univariate results in individually defined ROIs. Repeated-measures ANOVA with the factor NUMEROSITY and STIMULUS TYPE revealed significant main effects of numerosity in all ROIs (all $F(1,12) > 16.9$, all $p < 0.0001$). No main effect of stimulus type was observed in inferior and superior IPS (all $F < 0.69$, all $p > 0.42$); LOC showed a trend that did not pass the FDR correction threshold ($F(1,12) = 6.15$, $p = 0.029$). No interaction was observed in any of the ROIs (all $F < 1.3$, all $p > 0.28$). The results in inferior and superior IPS are similar to those obtained using literature-based ROIs (Figure 2). The numerosity effect in LOC differs from the (absent) numerosity effect reported in Figure 2. The difference might be due to the more dorsal location of individually defined LOC.

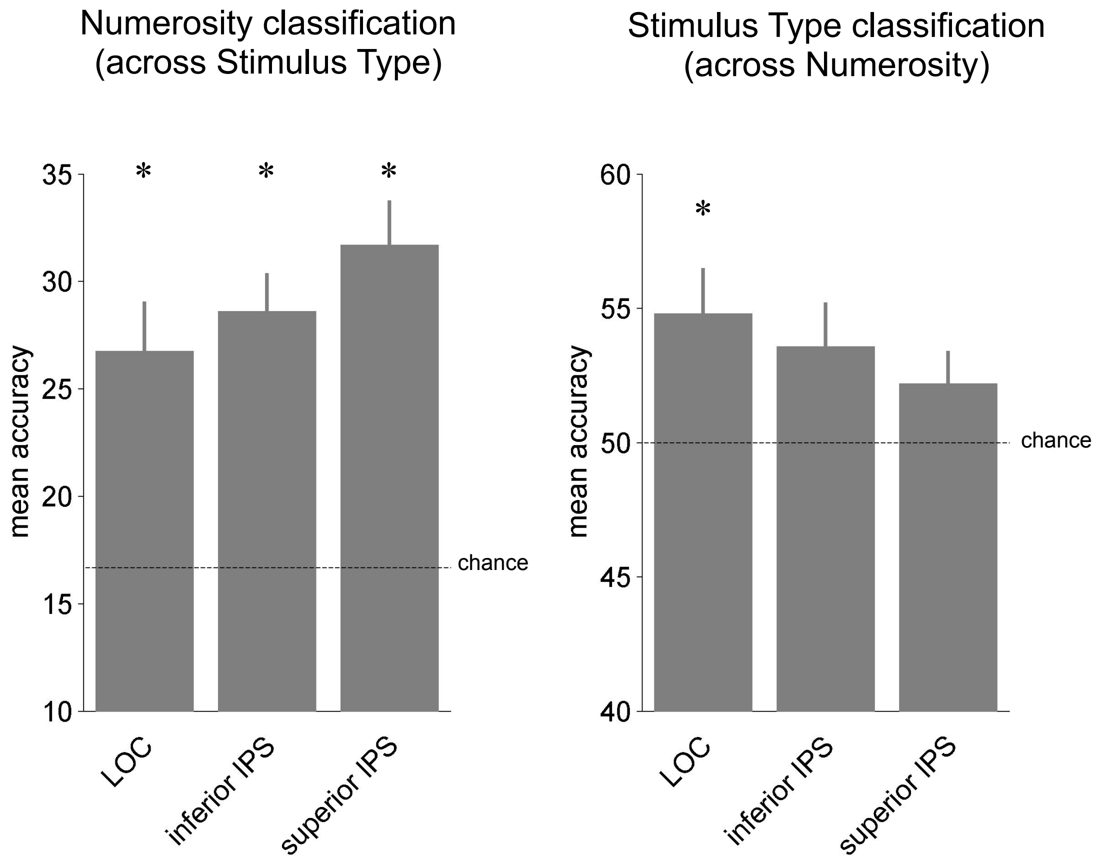


**Supplementary Figure 2.** Classification of Numerosity and Stimulus Type in individually defined ROIs. A repeated measures ANOVA with the factors ROI (LOC, inferior IPS, and superior IPS) and DECODING SCHEME (numerosity, stimulus type) revealed an interaction, indicating that decoding accuracies were highest for the numerosity decoding in inferior and superior IPS and for the stimulus type decoding in LOC ($F(2,22) = 3.9$, $p = 0.036$). In addition, a main effect was found for DECODING SCHEME, indicating higher decoding accuracies for numerosity decoding ($F(1,11) = 16.9$, $p = 0.002$). Paired t-tests did not reveal significant differences in classification strength between the ROIs after FDR correction (all $p > 0.22$). Higher decoding accuracies as compared to those revealed in literature-based ROIs (Figure 3) might be due to the subject-specific selection of peak regions in response to visual object processing (LOC, inferior IPS) and maintenance in short term memory (superior IPS) when using individually defined ROIs.

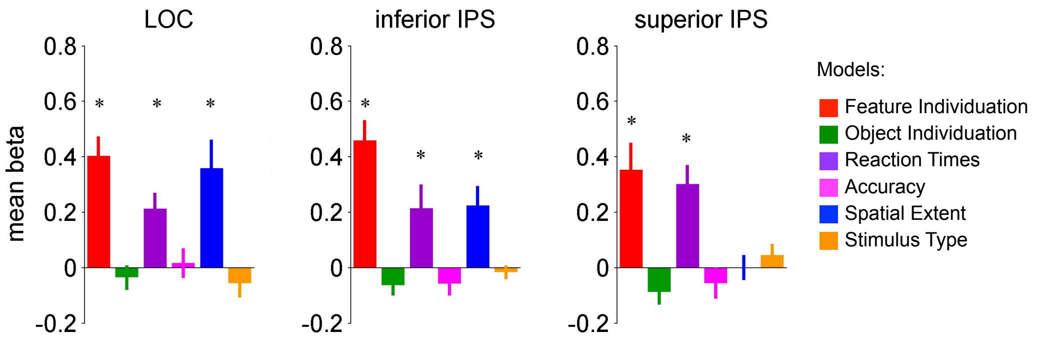


**Supplementary Figure 3.** Multiple regression RSA in individually defined ROIs. Asterisks indicate FDR corrected significant effects. Error bars indicate SEM. Paired t-tests between ROIs revealed significant differences for the Spatial Extent model in LOC vs. superior IPS ($t(12) > 3.45$, $p = 0.005$) and inferior vs. superior IPS ($t(11) > 3.87$, $p = 0.003$). The other models revealed no significant differences between ROIs.

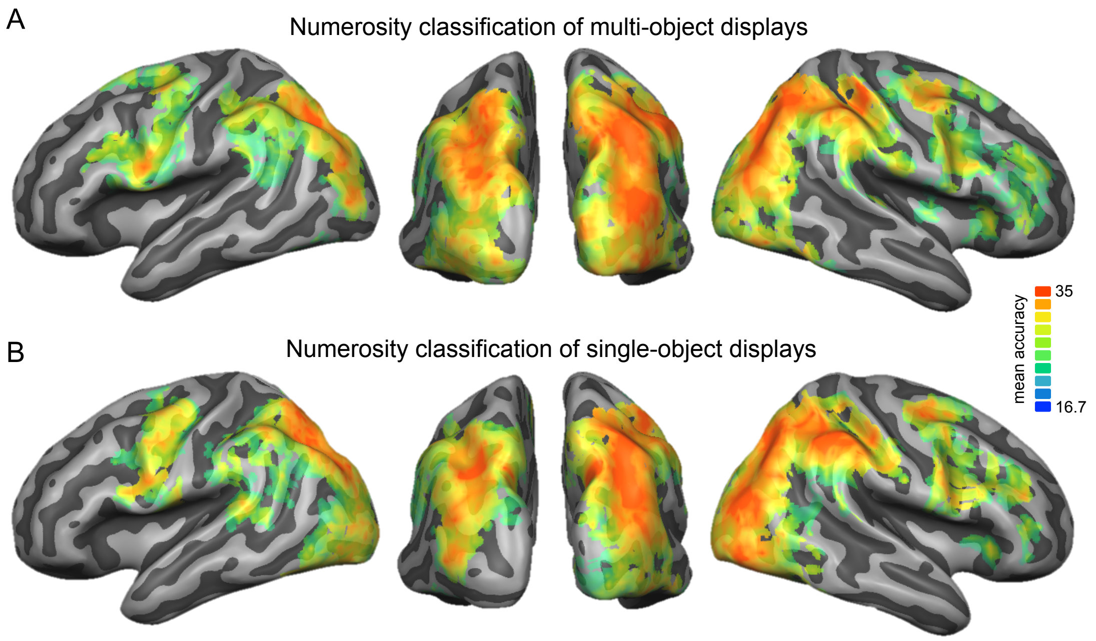


**Supplementary Figure 4.** Numerosity classification within multi-object (A) and single object (B) stimulus displays. Within each stimulus type, numerosities were decoded using leave-one-run-out cross validation, i.e., classifiers were trained to discriminate numerosities using data of 5 of 6 runs and tested on its accuracy to discriminate numerosities using data of the held out run (6 iterations). Accuracy for decoding at chance is 16.7%. Mean accuracy maps of the searchlight analysis were thresholded using cluster size correction for multiple comparisons at $p = 0.05$, initial voxel threshold $p = 0.001$.

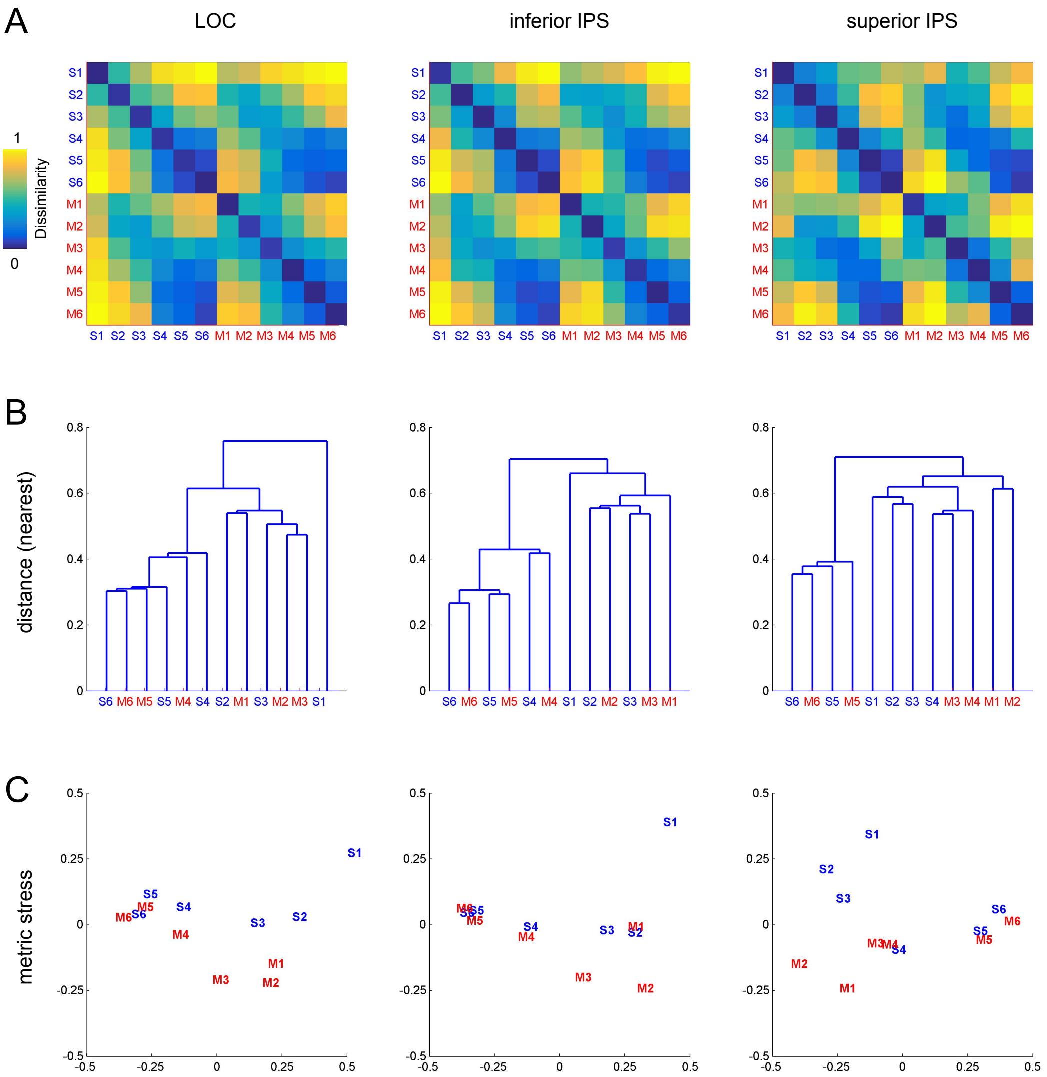


**Supplementary Figure 5.** Cluster analysis. (A) Representational dissimilarity matrices (1 – r, averages across participants) extracted from ROIs. (B) Dendrograms based on a cluster analysis (nearest distance). (C) Multidimensional scaling (metric stress). S1-S6 (in blue): single-object displays with 1-6 distinct target objects; M1-M6 (in red): multi-object displays with 1-6 parts of a single object.

**Supplementary Table 1**. Bayesian model comparison between different plateau models (plateau reached at numerosity of 4, 5, or not reached in the tested numerosity range, i.e., linear increase of BOLD response). Likelihood ratios between Bayes factors > 3 indicate moderate (and noteworthy) evidence in favor of one model vs. the other (Jeffreys, 1961).

| | plateau at 5 vs. plateau at 4 | | linear vs. plateau at 5 | |
|---|---|---|---|---|
| | single-object | multi-object | single-object | multi-object |
| inferior IPS | 136.90 | 21.85 | 4.81 | 2.29 |
| superior IPS | 31.89 | 18.68 | 2.39 | 1.62 |

**Supplementary Table 2.** Plateau analysis in ROIs using using individually defined ROIs. Results of paired t-tests investigating the point at which the increase in percent signal change for each additional target plateaued. Asterisks indicate comparisons that survive FDR correction.

| | LOC | | inferior IPS | | superior IPS | |
|---|---|---|---|---|---|---|
| | t(12) | p | t(12) | p | t(11) | p |
| *four vs. five targets* | | | | | | |
| single-object | -3.34 | 0.006* | -4.73 | <0.001* | -3.78 | 0.003* |
| multi-object | -2.14 | 0.054 | -2.82 | 0.015* | -4.68 | <0.001* |
| *five vs. six targets* | | | | | | |
| single-object | -0.80 | 0.44 | -1.31 | 0.216 | -2.66 | 0.022 |
| multi-object | -0.94 | 0.364 | -2.18 | 0.05 | -3.40 | 0.006* |

**Supplementary Table 3**. Bayesian model comparison between different plateau using individually defined ROIs.

| | plateau at 5 vs. plateau at 4 | | linear vs. plateau at 5 | |
|---|---|---|---|---|
| | single-object | multi-object | single-object | multi-object |
| LOC | 31.4 | 8.9 | 1.1 | 1.1 |
| inferior IPS | 462.4 | 35.2 | 5.5 | 2.9 |
| superior IPS | 220.3 | 207.6 | 7.7 | 8.3 |
